# Pore-modulating toxins exploit inherent slow inactivation to block K^+^ channels

**DOI:** 10.1101/576132

**Authors:** Izhar Karbat, Hagit Altman-Gueta, Shachar Fine, Tibor Szanto, Shelly Hamer-Rogotner, Orly Dym, Felix Frolow, Dalia Gordon, Gyorgy Panyi, Michael Gurevitz, Eitan Reuveny

**Affiliations:** Department of Biological Chemistry, Weizmann Institute of Science, Rehovot 76100, Israel; Department of Plant Molecular Biology and Ecology, Tel-Aviv University, Tel-Aviv 69978, Israel; Department of Biophysics and Cell Biology, University of Debrecen, 4032 Debrecen, Hungary; Structural Proteomic Unit, Weizmann Institute of Science, Rehovot 76100, Israel

## Abstract

Voltage dependent potassium channels (K_v_s) gate in response to changes in electrical membrane potential by coupling a voltage-sensing module with a K^+^-selective pore. Animal toxins targeting K_v_s are classified to “pore-blockers” that physically plug the ion conduction pathway and “gating modifiers” that disrupt voltage sensor movements. A third group of toxins blocks K^+^ conduction by an unknown mechanism via binding to the channel turrets. Here we show that Cs1, a peptide toxin isolated from cone snail venom, binds at the turrets of K_v_1.2 and targets a network of hydrogen bonds that govern water access to the peripheral cavities that surround the central pore. The resulting ectopic water flow triggers an asymmetric collapse of the pore by a process resembling that of inherent slow inactivation. Pore modulation by animal toxins exposes the peripheral cavity of K^+^ channels as a novel pharmacological target and provides a rational framework for drug design.

## Introduction

Voltage-gated potassium channels (K_v_s) shape the electrical properties of excitable cells by facilitating the selective flow of K^+^ ions across the cell membrane in response to changes in membrane potential. The K_v_ channel protein is composed of a pore domain that enable ion translocation, surrounded by four voltage sensing domains that gate the pore in a voltage dependent manner (Kuang et al., 2015). The selectivity for K^+^ is contributed by a 5-residue signature sequence (TVGYG) termed the “Selectivity Filter” (SF) that forms a constriction of the pore at the extracellular face of the channel (Heginbotham et al., 1994). To pass through the SF, K^+^ ions must shed their hydration shells. Having their backbone carbonyl oxygen atoms facing the pore and creating four consecutive coordination sites (s1-s4) for a dehydrated K^+^ ion, the signature residues catalyze this reaction (Zhou et al., 2001). At the SF, the alternate binding of K^+^ ions and water molecules in a single-file enables a “knock-on” mechanism, wherein an incoming ion exerts electrostatic repulsion that displace a neighboring ion along the pore, resulting in conduction (Roux et al., 2011).

In addition to its role in setting the preference for K^+^ ions, the SF serves as a gate that down-regulates ion flow via a process known as ‘C-type’ or ‘slow’ inactivation (SI). SI is triggered during prolonged depolarizations, and it is antagonized by ion occupancy in the pore (Cordero-Morales et al., 2006; Cuello et al., 2010; Yellen et al., 1994). While it is widely accepted that SI entails a conformational change that renders the SF non-conductive, the exact molecular description of this state is still under debate. The crystal structure of the bacterial channel KcsA at low external K^+^ has revealed a “pinched” SF in which the protein backbone was bent at Gly77 and ion coordination sites s2 and s3 were unoccupied (Zhou et al., 2001). This channel conformation was suggested to be sterically stabilized by a network of water molecules buried in cavities behind the SF (“peripheral cavities” hereafter, Ostmeyer et al., 2013). Residues situated near these cavities that serve as hydrogen-bond donors/acceptors were proposed to shape the rate of interconversion between the open and the inactivated channel states (Pless et al., 2013). This proposition is supported by the recent structure of the K_v_1.2–2.1 paddle chimera in lipid nanodiscs in which these hydrogen bonds are compromised. (Matthies et al., 2018)

Many venomous organisms carry in their arsenal peptide toxins that block K_v_ channels, thus lowering the threshold for an action potential at the affected nerve or muscle, leading to paralysis. These toxins have traditionally been classified into two groups based on their mode of action (Kalia et al., 2015). Gating modifier toxins block the channel by binding its voltage-sensing domain, thereby altering the stability of the closed state. Pore-blocker toxins bind at the pore module and physically occlude the ion permeation pathway. Classical K_v_ pore-blockers from scorpion venom are short polypeptides, typically 30-40 residue long, with a rigid core composed of an alpha helix and an anti-parallel 3-stranded beta sheet. Early experiments, in which the interactions of the scorpion neurotoxins Charybdotoxin (CTX) and Agitoxin2 with K_v_s were probed, revealed that a bound toxin can be dissociated by K^+^ ions entering the channel from the cytoplasmic side, a phenomenon termed “trans-enhanced dissociation” (MacKinnon and Miller, 1988). The latter phenomenon was completely abolished upon neutralization of the highly conserved Lys27 toxin residue (Hidalgo and MacKinnon, 1995; Park and Miller, 1992). A model in which the Lys27 ε-amino moiety of the bound toxin competes with K^+^ ions on a binding site at the extracellular face of the conduction pathway was put forward (Goldstein and Miller, 1993). This model was validated when the crystal structure of CTX in complex with a mammalian K_v_ channel was solved, revealing that the toxin projects the sidechain of Lys27 into the channel pore, and as a result K^+^ binding at the s1 site is lost (Banerjee et al., 2013).

In addition to these pore-blocking toxins, numerous studies have illuminated another diverse family of toxins that target the pore domain of K^+^ channels and block the ionic current, without physically occluding the pore. Instead, these toxins exhibit a pharmacophore made of a ring of positively charged residues and their binding sites are confined primarily to the channel turrets (Hu et al., 2014; Rodríguez De La Vega et al., 2003; Verdier et al., 2005; Xu et al., 2003; Zhang et al., 2003). The precise molecular mechanisms by which toxins of this family block their targets remain obscure, although a general concept (“turret-block”) whereby the toxin acts as a lid above the pore entry was proposed (Verdier et al., 2005; Xu et al., 2003).

Here we describe a novel block mechanism utilized by Conkunitzin-S1 (Cs1), a 60-residue peptide toxin isolated from *Conus Striatus*, which bind to the drosophila Shaker turrets. We show that instead of directly plugging the ion conduction pathway, Cs1 modifies the permeation of water molecules into the peripheral cavities, thus creating highly asymmetric water distributions around the SF that trigger a local collapse of the channel pore, analogous to slow inactivation. In addition to the description of the “pore-modulatory” action of the toxin, the data provides general new insights into the usage of the peripheral cavities of potassium channels as an important therapeutic target and a framework for rational drug design to affect channel function.

## Results

### Molecular dissection of the Conckunitzin-S1 – Shaker_KD_ complex

Conckunitzin-S1 (Cs1) is a 60-residue toxin isolated from the venom of the fish hunting cone snail *Conus striatus* (Bayrhuber et al., 2005). The toxin molecule has a conserved Kunitz domain fold, composed of an N-terminal 3-10 helix, two-stranded beta sheet and a C-terminal alpha helix reticulated by two disulfide bridges (Dy et al., 2006). Our initial analysis of the toxin has indicated that it is highly toxic to fish larvae (*Danio rario*, LD_50_=500nM in bath application), practically non-toxic to mice (LD_50_ > 50μg/gr by subcutaneous injection), and that it has a considerable preference for insect over mammalian K_v_ channel isoforms (Fig. S1A, see also Schmidt et al., 2014). Aimed to decipher the structural basis for this preference, we carried a molecular dissection of the toxin-channel complex. First, the solvent exposed residues of Cs1 were substituted by alanine, the resulting toxin mutants were expressed and purified, and their potency on the high affinity K427D mutant of the drosophila *shaker* channel (dmK_v_1.2 K427D, Shaker_KD_ hereafter) expressed in *Xenopus* oocytes was assayed. This analysis has pinpointed 5 toxin residues important for toxicity, four of which were confined to the C-terminal α-helix (Fig S1B). Site directed mutagenesis targeted to the extracellular residues of the channel pore domain revealed three residues with critical contributions to Cs1 binding – the aromatic Phe425, and the nearby negatively charged Asp427 and Asp431 (Fig. S1E,F). We concluded the molecular dissection with a double-mutant cycle analysis (Hidalgo and MacKinnon, 1995; Horovitz, 1996) to expose putative toxin-channel amino acid pairwise interactions (Fig. 1C). This analysis revealed substantial coupling energies between Arg34 of the toxin and all three channel residues, as well as between Phe425 of the channel to Arg49 and Tyr59 of the toxin. We set to obtain putative structures of the channel-toxin complex by employing a complementary, independent approach of non-constrained rigid-body docking. Toward this aim we crystalized and determined the structures of two toxin mutants at 1.3Å resolution (Fig S1D, Materials and Methods) from which we obtained high quality model of Cs1. The toxin molecule was docked onto a channel model that was based on the 2.4Å crystal structure of the closely-related k_v_1.2-k_v_2.1 paddle chimera (Long et al., 2007). Clustering the obtained putative structures, we observed a dominant toxin binding mode that accounted for 836 of the top 1000 solutions. This model was compatible with the distance restrains provided by the experimental coupling energies (Fig 1B), and it was stable during prolonged molecular dynamics (MD) simulations (>0.4μs). The docked toxin model revealed a network of hydrogen bonds and salt bridges between the side chains of Shaker_KD_:Asp447 from three of the four channel domains and the C-terminal residues of the toxin (Fig 1 C,D). These interactions could not be deduced by molecular dissection since substitutions at Shaker_KD_:Asp 447 which resides at the SF entrance, results in a non-conductive channel mutants (Yifrach and MacKinnon, 2002, and not shown). The positively-charged side chain of Cs1:Arg34 interacts with a negative binding pocket formed by Shaker_KD_: Asp427 and Asp431, rationalizing the weak binding of the toxin to the wild-type channel bearing a lysine at position 427 (Fig. S1A). The structural basis for the preference of Cs1 for insect over mammalian K_v_ channels can be readily inferred – the side chain of Phe425 from channel subunits B, C and D makes high impact interactions with Tyr59, Arg49 and Arg34 of Cs1, respectively. The high diversity of these interactions – π− π, cation- π, and aromatic-aliphatic stacking interactions mandates an aromatic residue at this position for high affinity binding. This criterion is frequently met by K_V_1.2 isoforms isolated from crustaceans, fish and insects, but it is not observed in mammals (Fig. S1G), consistent with the preferential toxin action on the natural prey of the cone snail.

**Figure 1:**
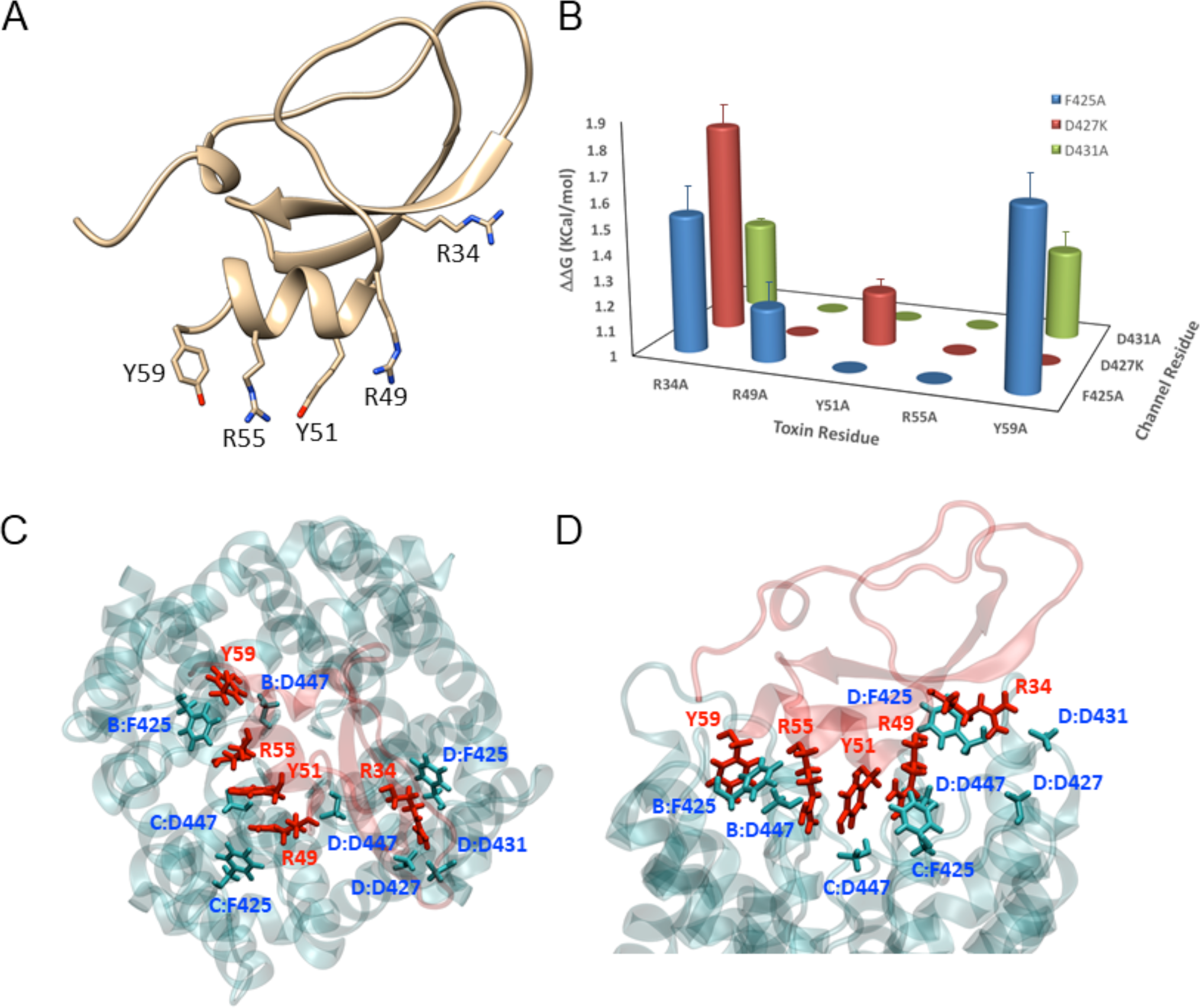
Molecular dissection of the Conckunitzin-S1 – Shaker_KD_ complex. **A**, The biocative residues of Cs1. The presented structure was extracted from a snapshot taken following a 10ns equilibration run of the Shaker_KD_ - Cs1 complex (see methods). Toxin backbone is rendered in ribbons, residues whose substitution by alanine decreased free energy by >1 Kcal/mol (Fig. S1B) are rendered in sticks. **B**, Double-mutant cycle analysis of Shaker_KD_ - Cs1 interactions. The absolute ΔΔG values obtained by assaying the potency of the indicated toxin mutants on each of the indicated channel mutants are shown. The data were thresholded at 1 Kcal/mol to isolate the high-impact interactions. **C, D**, MD-equilibrated structure of the Shaker_KD_-Cs1 complex, top (C) and side (D) views. Channel and toxin molecules are rendered in blue and red transparent ribbons, respectively. Interacting residues are rendered in sticks. Channel residues are indicated using a chain:residue notation.

### Conkunitzin-S1 does not directly block the ion conduction pathway

A distinctive feature of the modeled Cs1-Shaker_KD_ complex that became evident during early MD refinements was that the toxin does not physically occlude the channel pore. Cs1 is bound slightly off the central pore axis, and none of the toxin residues interact with the SF backbone carbonyls, as do classical K_v_ pore blockers (Goldstein and Miller, 1993). In MD simulations conducted with bound Cs1, water exchange between the outer vestibule of the channel pore and the bulk was observed, albeit at reduced rate compared to a toxin-free channel (Fig 2A,B). The gap between the toxin molecule and the channel pore was often occupied by a fully hydrated K^+^ ion (movie S2). In stark contrast, MD simulations of the classical pore blockers CTX and ShK (Lanigan et al., 2002) bound to their respective receptors, revealed a tight seal formed by the toxins around the channel pore (Fig. 2A,B, movie S2). This disparity between the simulated behavior of Cs1 and the canonical pore blockers lends itself to experimental scrutiny using trans-enhanced dissociation experiments (MacKinnon and Miller, 1988). In these experiments, we assayed toxin dissociation during strong depolarizations, which drive K^+^ ions to compete with a bound toxin on the S1 ion coordination site. We have elected to utilize for these experiments the classical pore-blocker ShK and not the wildly known Charybdotoxin, for which a complex structure is available. The reason for our choice was that CTX does not tolerate well an aromatic residue at position 425 of the Shaker channel (Goldstein and Miller, 1992), while Cs1 mandates it (present work). In contrast, ShK is quite tolerant to substitutions at position 425 (Fig. 5B). While ShK could be readily knocked off its binding site by prolonged depolarizations, Cs1 remained bound to the channel throughout these experiments, consistent with the proposed docking model in which none of Cs1 residues is bound at the s1 site (Fig 2C,D).

**Figure 2:**
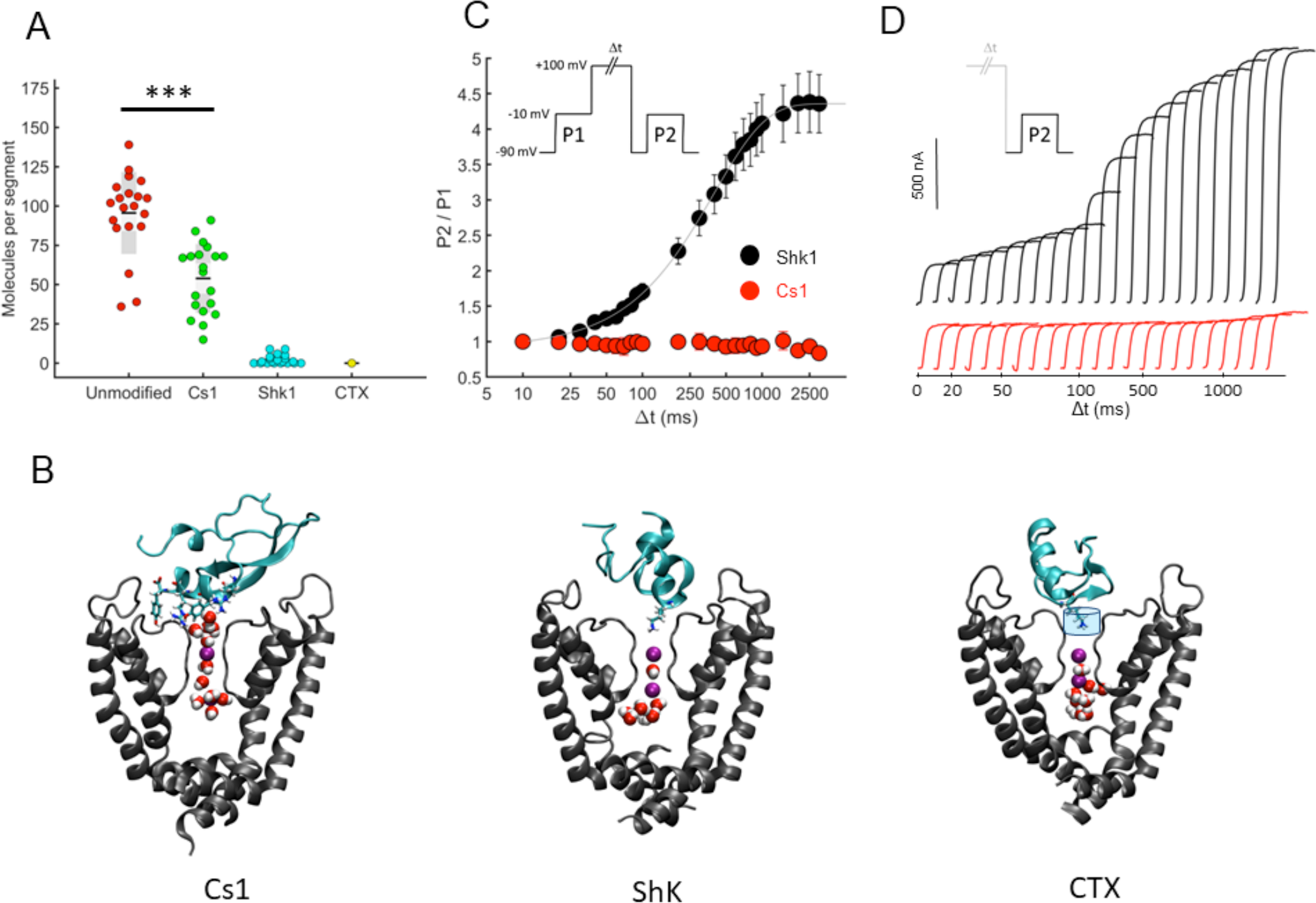
Conkunitzin-S1 does not directly block the ion conduction pathway. **A**, The effect of bound toxins on water traffic at the extracellular vestibule of Shaker_KD_. 200ns MD trajectories of unmodified channel, or of channel bound to each of the indicated toxins, were segmented into 20 X 10ns windows. At each segment, the number of water molecules passing through a cylindrical volume positioned just above the SF (depicted in panel B, CTX) were counted. In unmodified Shaker_KD_, an average of 95.6±26 water molecules per segment were observed, compared to 53.95±21.88, 2.2±2.9 and 0 molecules per segment for Cs1, ShK and CTX, respectively. **B**, Cross sections through the protein channel complex, taken at the end of a 20ns equilibration MD run, revealing the water and ion occupancy near the pore. Two opposite channel subunits (gray) bound to a toxin molecule (cyan) are depicted. the bioactive residues at the C-terminus of Cs1 as well as Lys22 and Lys27, of ShK and CTX, respectively, are shown. K^+^ ions and water molecules residing near the pore are rendered in space-fill. A cylinder of 2Å radius and 3Å in height positioned 1Å above the plane of outermost SF residue (Gly446), used for water traffic measurements (A), is overlaid on the CTX complex structure. **C**, Cs1 is insensitive to trans-enhanced dissociation. Toxin dissociation was induced using the voltage protocol depicted at the inset as described in Materials and Methods. The effect of ShK was completely reversed by prolonged depolarizations with a single exponential time-course (*τ* = 405±51 ms, max = 4.4±0.77, n = 4), while Cs1 has remained bound throughout the experiment (P2/P1~1). **D.** Typical currents recorded during P2 as a function of Δt in the presence of Cs1 (red) or ShK (black). P2 Currents from individual traces are overlaid using an arbitrary horizontal displacement, Δt is the length of the depolarizing pulse preceding P2.

### Conkunitzin-S1 induces an asymmetric constriction of the selectivity filter

The results presented thus far strongly negate a physical block of the channel pore by Cs1 and suggest that water molecules and ions from the extracellular milieu may access the outer channel vestibule in the presence of a bound toxin. Yet, ion flow is blocked completely when a saturating toxin concentration is applied (Fig S1A). To resolve this apparent conundrum, we simulated the behavior of the toxin bound channel during a series of 200ns time windows. In 5 out of 7 simulations carried, we observed a non-symmetric collapse of the channel pore at the SF region (Fig. 3). The observed collapse occurred typically after few tens of nanoseconds into the production run, and followed a common pattern of molecular events, depicted in Figure 3. In all simulations, the SF was initially found at 2,4 ionic configuration, with a K^+^ ion bound at s2 and additional ion fluctuating between s4 and the intracellular cavity of the channel (Fig 3B). At this stable ionic configuration (in the absence of an electric field), the SF assumed a symmetric conformation with a cross-subunit distance of ~8Å (Fig 3C). The initial trigger for pore collapse was the flip of the backbone carbonyl at Gly444 or Tyr445 away from the pore into the peripheral cavity (Fig 3A). This event was closely followed by the dissociation of the K^+^ ion bound at s2, leaving the SF with a single ion occupying s4, and water molecules at the remaining sites (Fig. 3B). This configuration of the SF was highly unstable, and triggered a rapid asymmetric collapse of the channel pore, in which one pair of opposing subunits assumed a pinched conformation typified by a 5.5Å cross-subunit distance (Fig. 3C). This phenomenon was unique to Cs1 simulations, and was not observed in control simulations using either toxin-free channels (Fig 3), that were obtained using the very same starting configuration with the toxin molecule removed, or complexes of the classical pore blockers, CTX and ShK, bound to K_v_ channels (Fig. S3). On the contrary, in the latter case the constant occupancy of ion coordination site s1 by the ε-amino group of the conserved toxin lysine stabilized the SF conformation, manifested in decreased fluctuations of the backbone atoms and the bound ions (Fig 3, S3). These observations strongly suggest that the asymmetric collapse observed at the selectivity filter region in our simulations is linked to the presence of the bound toxin rather to the initial configuration of the system.

**Figure 3:**
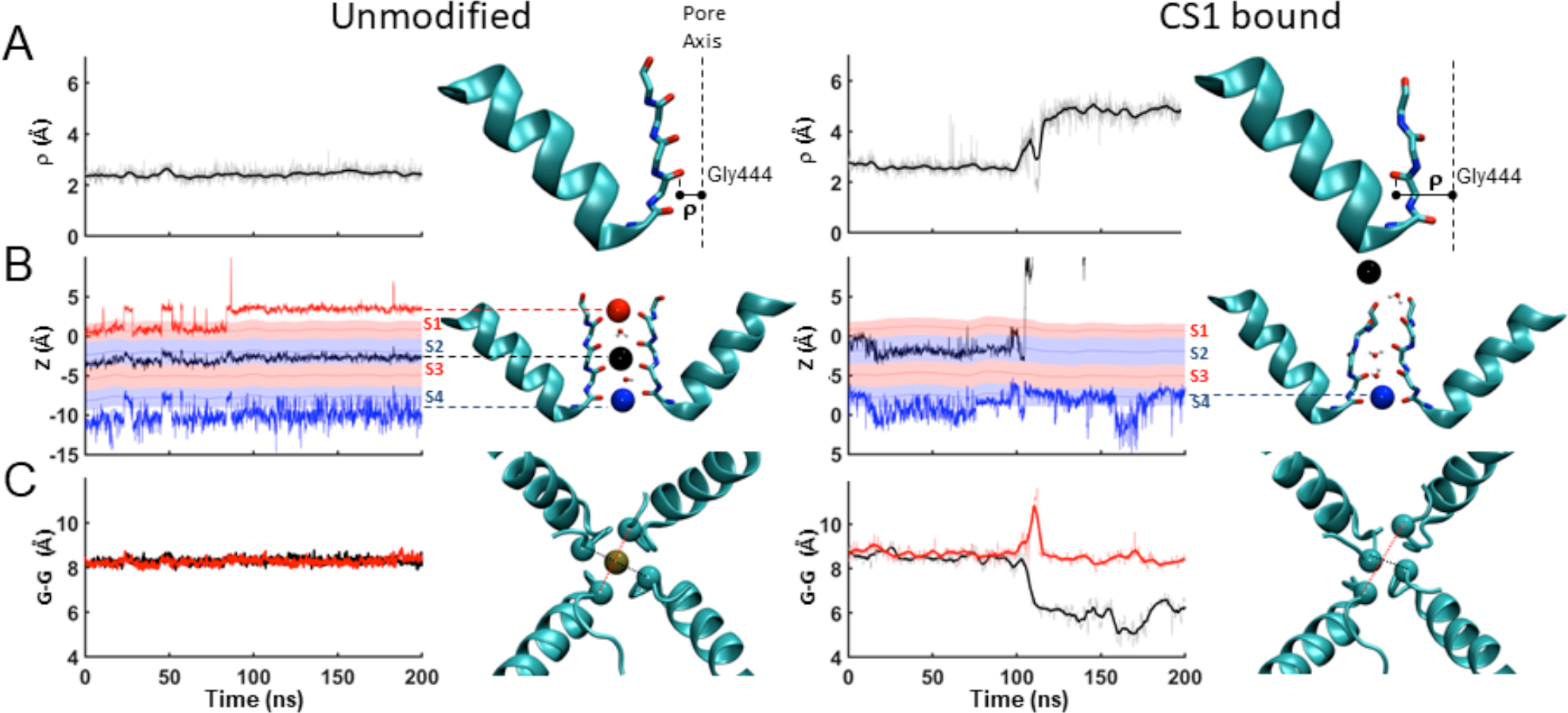
Conkunitzin-S1 induces an asymmetric constriction of the selectivity filter. The integrity of the selectivity filter region during 200ns long trajectories of unmodified (left) and CS1-bound (right) Shaker_KD_ channels. Drawings to the right of each time series depict the monitored parameters at the 200ns time-point. **A**, Flipping of the backbone carbonyl atom of Gly444(**G**YG). The radius of gyration (ρ) is a measure of the distance between the oxygen atom and the geometrical center of the pore. In the unmodified channel ρ is kept at 2.41±0.19Å throughout the simulation, in the toxin bound channel it has increased from 2.62±0.08Å (0-50ns) to 4.8±0.11Å (150-200ns). **B**, The vertical coordinates (Z) of K^+^ ions in the SF along the MD trajectory. Alternating red-blue stripes at the background delineate the s1-s4 ion coordination sites. The drawing depicts the SF region of two diagonally opposed subunits. Ions are rendered as solid spheres and colored to match the time-series plot. Pore-resident water molecules are rendered in CPK. Three Ions remain bound to the SF of the unmodified channel throughout the trajectory, whereas an ion originally coordinated at s2 is lost in the toxin-bound channel at t~100ns. **C**, Cross-subunit distance between the Cα atoms of Gly444 residues from chains A and C (red) or B and D (black). This distance is kept at 8.3±0.14Å during the unmodified channel simulation. During the toxin-bound simulation, an asymmetric constriction of the pore is observed following the dissociation of the ion from s2 (b), as GG (chain B: chain D) is decrease from 8.59±0.24Å (0-50ns) to 5.8±0.6Å (150-200ns), while GG (chain A: Chain C) remains roughly unchanged. The drawing provides a top view of the channel pore, the Cα atoms of Gly444 residues are rendered in space-fill and the GG distances are indicated.

### Conkunitzin-S1 modifies water permeation into the peripheral cavities

The observed collapse of the channel pore in the presence of Cs1 was initiated by a flip of a backbone carbonyl at Gly444 (Fig. 3A). This pore residue is positioned ~11Å away from the nearest toxin residue (Arg55) and ~15Å away from the channel turrets, where all high impact interactions with the toxin take place. How can Cs1 remotely trigger a cascade of events that leads to the collapse of the pore? A long-range allosteric effect mediated by the turret residues, which contribute most of the Cs1 binding site, seemed unlikely since (i) substitutions at these residues had no apparent effect on the stability of the open state (Ranganathan et al., 1996; Yifrach and MacKinnon, 2002) and (ii), comparison of the conformational dynamics of the channel backbone during system equilibration phase with and without bound toxin did not reveal any signs for induced fit (backbone RMSD between the initial and the equilibrated structures were 1.9±0.07Å and 1.93±0.16Å (n=5) for toxin free and toxin bound simulations, respectively). We therefore focused on the interactions between the bound toxin and channel residues in the vicinity of the pore region. The central pore of K^+^ channels is surrounded by water-filled cavities, which were proposed to shape the rate and extent of the slow inactivation process (Ostmeyer et al., 2013). In K_v_1.2, the confinement of water molecules within these peripheral cavities is achieved by two sets of channel residues. The “aromatic cuff” (Pless et al., 2013, Fig. S4A), composed of the side chains of Tyr445, Trp434 and Trp435 alongside Val443 form a hydrophobic barrier at the base of the cavity (“lower barrier”). At the top of the cavity, the side chains of Asp447, Met448 and Trp434 from one channel subunit and Thr449 from a neighboring subunit form a barrier that limits the exchange of water molecules between the cavity interior and the extracellular bulk (“upper barrier”, Fig S4B, movie S4A). In simulations of toxin free channels, the space between the two barriers is stably populated by 2-3 water molecules that exchange with the bulk at a typical average rate of ~30 molecules per 100ns (Fig. S4G). Since this exchange involves the crossing of the upper barrier, its rate depends on barrier dynamics. In particular, we find high correlation between the rate of exchange of cavity water and the dynamics of the hydrogen bond formed between the side chains of Asp447 and Trp434 (Fig. S4G), which was previously dubbed a “molecular timer” that sets the pace of slow inactivation (Pless et al., 2013, Fig. S4A). Since we find a strong correlation between its integrity to the rate of water exchange via the upper barrier, we refer to it hereafter as the D447:W434 gate, or simply the “D-W gate”. Analysis of Shaker_KD_ trajectories in the presence of Cs1, reveal a major impact of the toxin on the dynamic behavior of the D-W gate. Asp447, positioned directly above the selectivity filter, interacts electrostatically with toxin residues in three of the four channel subunits (Fig.1, Fig 4A, S4D). These contacts allow the toxin to modify the D-W gate in a non-symmetrical fashion. In channel subunit C, a hydrogen bond and a salt bridge between Asp447 to Tyr51 and Arg55 of the toxin, respectively, stabilize the D447:W434 hydrogen bond, leading to a “closed” conformation of the D-W gate (Fig. 4B). Conversely, in subunit D of the channel, a salt bridge between Cs1:Arg49 and Asp447 “pulls” the later away from Trp434, leading to a constantly “open” gate conformation (Fig. 4B, S4H). In addition, we observed strong effects exerted by the interactions of Cs1:Tyr59 with the upper barrier residues in channel subunit B (Fig S4D, movie S4C) on the permeation of water into the peripheral cavity.

**Figure 4:**
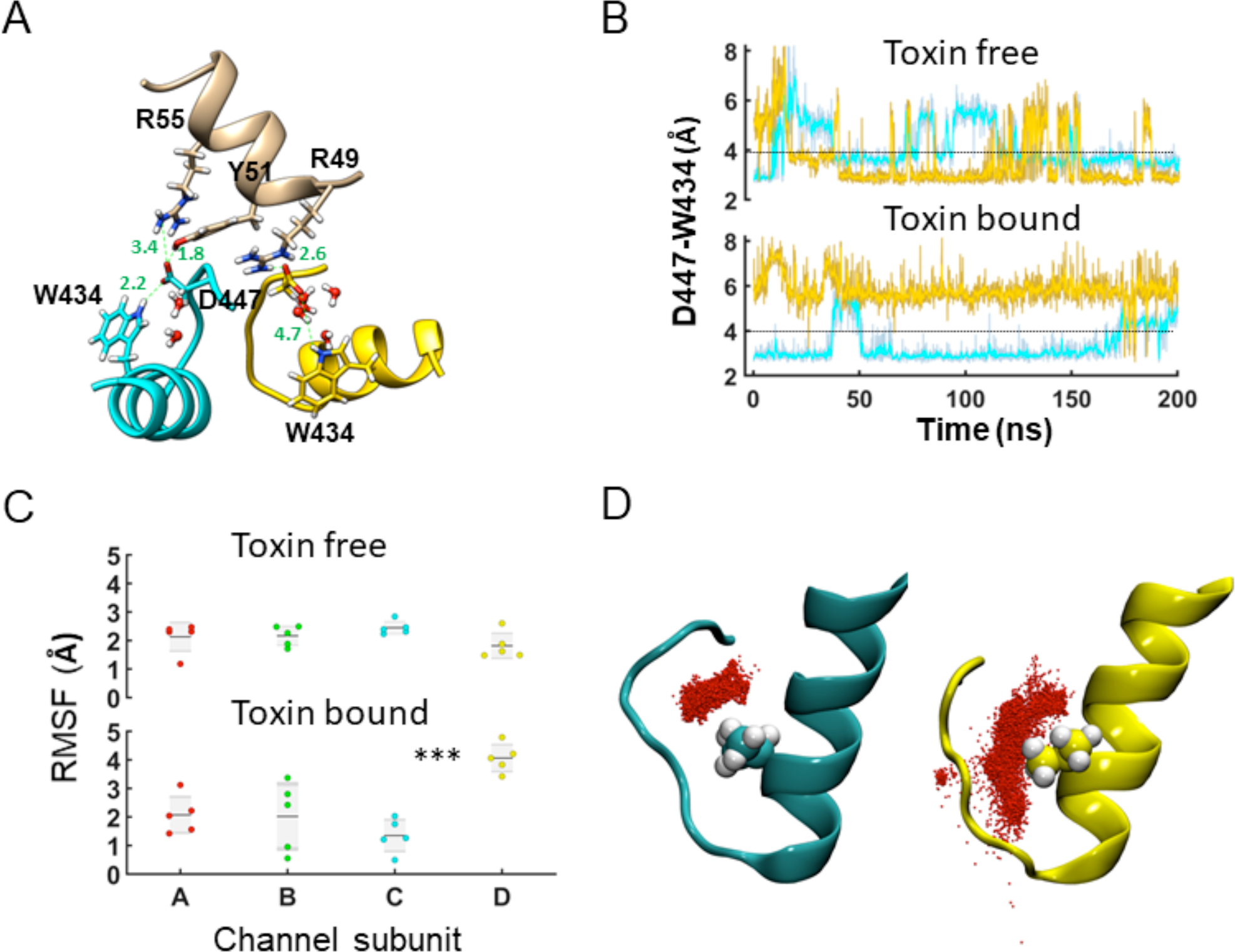
Conkunitzin-S1 modifies water permeation into the peripheral cavities. **A**, ribbon diagram depicting the key interactions between the alpha helical segment of the toxin (tan) and the Asp447 sidechains of subunits C (cyan) and D (gold). Green labels indicate distances in angstroms. **B**, conformational dynamics of the W434-D447 bond in subunits C (cyan) and D (gold) of unmodified and toxin-bound Shaker_KD_. The distance between the O_δ_1/O_δ_2 atoms of Asp447 and the N_ε_1 atom of Trp434 along a 200ns trajectory is plotted, a broken line marks the cutoff distance for a hydrogen-bond. **C**, the root-mean-square fluctuations (RMSF) of water molecules inside the peripheral cavities of unmodified and toxin-bound Shaker_KD_. Each data point represents the mean RMSF over an independent 200ns production run. **D**, spatial distribution of water molecules within the peripheral cavities of subunits C (cyan) and D (gold) over a 200ns trajectory. Dots mark the centers of the water oxygen atoms. The sidechain of Val438 positioned at the base of the aromatic cuff barrier is rendered in spheres.

**Figure 5:**
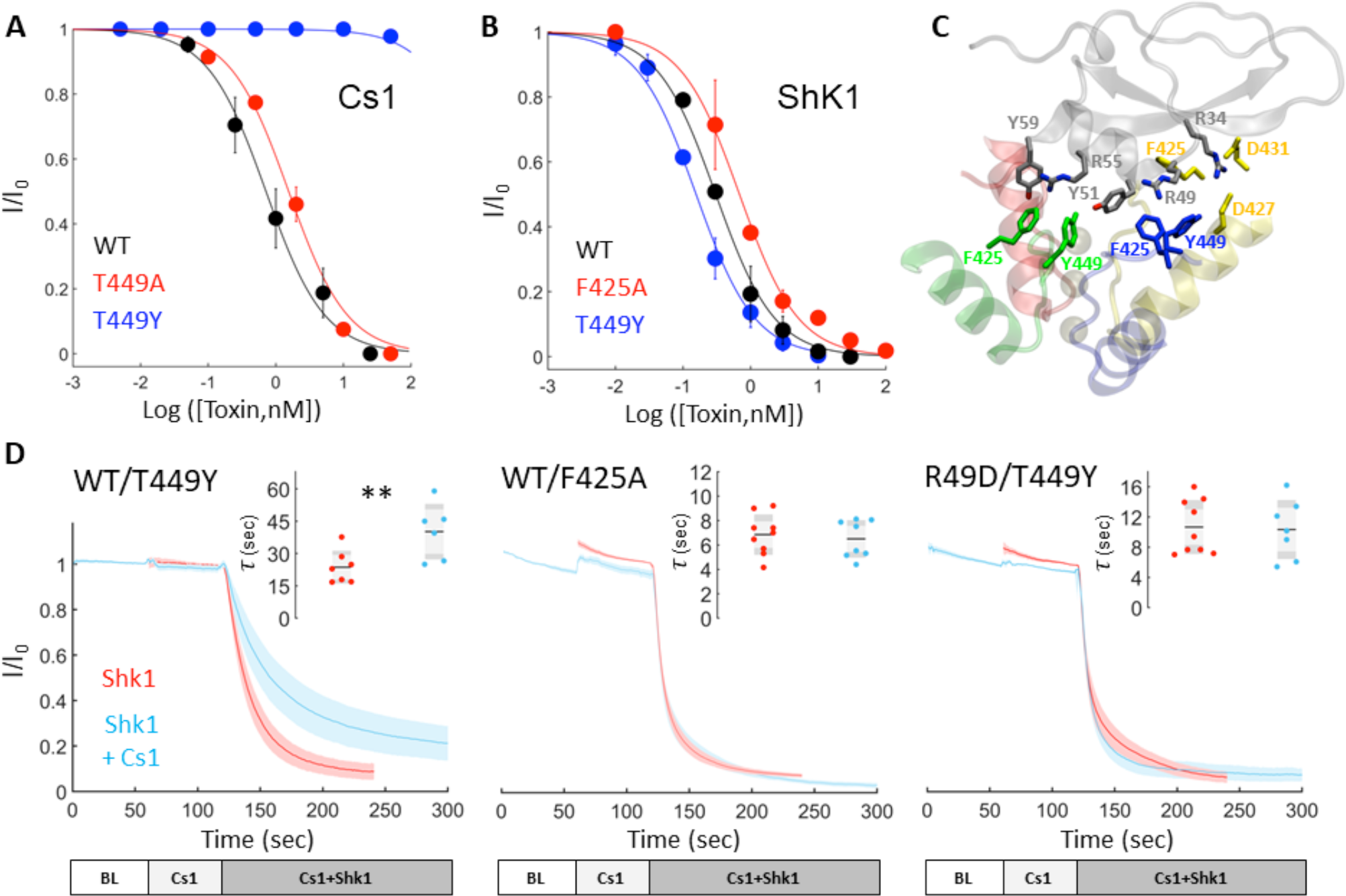
A silent binding mode of Conkunitzin-S1 to a non-inactivating Shaker mutant. **A, B**, Dose-response curves of Cs1 (A) or ShK (B) determined on Shaker_KD_ derivatives expressed in oocytes. The EC_50_ values obtained are 0.95±0.03 nM (unmodified), 1.92±0.64 (T449A) and >1μM (T449Y) for Cs1 (A), and 0.29±0.11nM (unmodified), 0.68±0.28nM (F425A), 0.16±0.06 (T449Y) for ShK (B). **C**, snapshot of the Cs1-Shaker_KD_ T449Y complex taken following 200ns unconstrained MD simulation. Interacting residues are rendered as sticks, channel pore-helices and the toxin are rendered as transparent ribbons. This structure retains all the high impact toxin-channel interactions identified by double-mutant cycle analysis (Fig. 1B). Note the newly introduced sets of cation-*π* and *π*-*π* interactions between the tyrosine substituted at position 449 and the C-terminal residues of the toxin. **D**, Kinetics of Shaker_KD_ block by ShK alone or in the presence of Cs1. Peak K^+^ currents were recorded from oocytes expressing Shaker_KD_ T449Y (left, right) or F425A (middle) for 60 seconds, and then 5nM ShK were applied to the bath and the resulting current decay was recorded (red traces). In a coupled set of experiments conducted on oocytes from the same batch (blue traces), the introduction of ShK was preceded by the application of 50nM Cs1 (left, middle) or 50nM of the weakly active (EC_50_ = 66.4±5.4nM) Cs1 R49D mutant (right). Bars at the bottom indicate the timing of toxin application (BL – baseline). Shaded area behind the curves represents standard deviation. Decay time constants calculated with (blue) and without (red) pre-application of Cs1 are given at the insets.

### Asymmetric water permeation into the peripheral cavities promotes pore-collapse

The asymmetric effect of Cs1 on the open probability of the D-W gate in discrete channel subunits directly translated in MD simulations into an asymmetric rate of water exchange through the upper barriers. In toxin-free simulations, the rate of water exchange between the peripheral cavities and the bulk solvent through the upper cavity barriers was 31.75±2.4 molecules per 100ns, evenly distributed between the four channel subunits (Fig S4E). In Cs1-bound simulations, the rate of water exchange through the upper barrier in subunit D, having a permanently open D-W gate, was nearly four-fold higher compared to subunits B and C, having their D-W gates in predominantly closed conformations (Fig S4E, S4H). This resulted in a high thermal agitation of water molecules within the peripheral cavity of subunit D, compared to the other subunits (Fig 4C). These highly disordered water molecules could not be stably confined within the peripheral cavity boundaries, and instead they spread to neighboring channel regions by triggering a flip of the aromatic ring of Tyr445, thereby breaching the hydrophobic barrier at the bottom of the cavity (Fig. 4D, movies S4B,S4C). The breach of the lower barrier allowed the exchange of water with the neighboring cavities (movie S4A, Fig. S4G), or with the intracellular vestibule through a path contributed by the hydrophilic side chains underneath (movie S4B,S4C, Fig S4F). In both routs, the flow of water behind the selectivity filter could be diverted into the central pore by a flip of a backbone carbonyl at Gly444 or Tyr445, displacing the bound ion from S2 and leading to pore collapse (Fig 3, movie S4C). In turn, collapse of the central pore increased the diameter of the peripheral cavity, allowing for faster water flow behind the selectivity filter into the intracellular vestibule of the channel pore (movie S4C).

### A silent binding mode of Conkunitzin-S1 to a non-inactivating Shaker_KD_ mutant

The described mechanism of Cs1 action is highly reminiscent of the inherent slow inactivation, as both processes involve the constriction of the channel pore in response to alternating hydration patterns at the peripheral cavities. A mutation at the peripheral cavity upper barrier, T449Y, previously shown to eliminate slow inactivation (López-Barneo et al., 1993) also abolished block by Cs1 (Fig. 5A). Yet, we did not assign this residue to the Cs1 binding site, as both mutagenesis (Fig. 5A) and modeling data did not reveal any contribution of Thr449 for toxin binding. The finding that the T449Y mutation has not affected the binding of ShK (Fig 5B), which makes close contacts with the pore region, suggested that the substitution has not induced major rearrangements within the channel protein. To gain structural insight into the loss of function of Cs1 on Shaker_KD_ T449Y, we introduced the mutation into the docked toxin model and subjected the resulting complex to MD simulations. This complex retained all high impact toxin-channel contacts and exhibited novel favorable interactions between the toxin and the substituted tyrosine (Fig. 5C). Simulation therefore predicts a tight binding of Cs1 to the Shaker_KD_ T449Y mutant, despite its apparent lack of activity. We tested this prediction by a series of functional binding competition assays in which the ability of Cs1 to inhibit the binding of ShK to Shaker_KD_ derivatives expressed in oocytes was examined (Fig. 5D). Indeed, we found that the rate of ShK association with Shaker_KD_ T449Y was reduced upon pre-application of Cs1 (Fig. 5D, Fig S5A,B). This effect was specific for this channel mutant - we have not observed competition between the two toxins on a channel mutant with low affinity for Cs1 (F425A, Fig 5D, middle), or upon pre-application of a toxin mutant with low affinity for the channel (R49D. Fig 5D, right). How could the apparent high-affinity binding of Cs1 to Shaker_KD_ T449Y be reconciled with its apparent lack of activity? MD simulations of the Cs1-bound T449Y channel mutant revealed high stability of the channel pore (Fig. S5C). Analysis of these trajectories offered two non-redundant mechanisms (Fig. S5) that rationalize the pore stability in the presence of bound toxin. The first is the stabilization of the aromatic-cuff barrier by the newly introduced tyrosine rings. In subunits A and D of the channel, the Tyr449 rings rotated downwards and formed multiple contacts with residues of the aromatic-cuff barrier. This has stabilized the barrier and allowed it to resist the highly disordered water molecules allowed into the peripheral cavities by the bound toxin (Fig. S5C, movie S5A). A second mechanism involved a structural rearrangement within the channel-toxin complex, in which the critical interactions between the C-terminal residues of the toxin and the D-W gates at chains C and D were replaced by a novel set of interactions with the newly introduced tyrosine rings (Fig. S5C-E, movie S5B). This rearrangement has diminished toxin effect on D-W gate dynamics and restored normal water permeation into the peripheral cavities. In summary, the silent binding of Cs1 to Shaker_KD_ T449Y strongly support our supposition that the toxin does not directly plug the ion conduction pathway, and simulations of the channel mutant with a bound toxin reveal specific modifications focused at the proposed action sites of the toxin.

### Channel block by Conkunitzin-S1 is slowed by heavy water

The proposed blocking mechanism for Cs1 postulates two stages of toxin action. The first is channel binding, driven mainly by aromatic and cation-π interactions between the toxin and the channel turrets. This stage by itself does not block the channel pore. The second is the interference of the toxin with the normal behavior of the D-W gates via polar interactions, which triggers pore-collapse. This second step crucially depends on the flow of water molecules through the peripheral cavities that surround the selectivity filter. A prediction derived from the proposed mechanism would be that the rate of channel block by the toxin is a function of the basal exchange rate of water molecules between the cavity interior and the bulk solvent. To examine the validity of this prediction, we have sought an experimental design that would allow us to separate between the two steps of Cs1 action, namely toxin binding and pore-collapse. To this end, we have employed the Shaker_KD_ M448K mutant, which exhibit accelerated slow inactivation, while retaining high affinity for toxins (Fig S6A; Koch et al., 2004). While Cs1 binding to unmodified Shaker_KD_ was voltage-insensitive (Fig. 2C,D), we could partially reverse toxin block of the M448K mutant using prolonged depolarizations (Fig. S6B). This voltage-induced reversal of channel block was distinct from classical trans-enhanced dissociation (Fig. S6B). The toxin effect could be restored by the subsequent incubation at a potential where the channel is closed, at a rate that was toxin concentration dependent, giving rise to a bell-shaped curve, with a rising phase that follows recovery from slow inactivation, and falling phase that follows re-establishment of toxin block (Fig 6A, S6C). The rate constants associated with these two opposing processes typically differed by an order of magnitude allowing for a straightforward isolation of Cs1 block onset rate with high temporal resolution (Fig.6). To further isolate the rate component contributed by the water exchange at the peripheral cavities, a series of experiments in which we compared the function of the toxin in H_2_O and D_2_O based solutions was performed. D_2_O was chosen for these experiments since, (*i*) it is compatible with electrophysiological measurements; (*ii*), it has a smaller diffusion coefficient compared to H_2_O (Weingärtner, 1984), and (*iii*), deuterium bonds are more stable compared to hydrogen bonds (Scheiner and Čuma, 1996). We reasoned that the latter two factors should slow down the D_2_O-exchange rate through the upper barrier and allow us to challenge directly the proposed action mechanism of Cs1. Since the effect of D_2_O on the rate of cavity water exchange in Shaker_KD_ is not readily amenable for direct experimental determination, we sought to examine it indirectly, taking advantage on the established link between the water occupancy at the peripheral cavities and the macroscopic rate of recovery from slow inactivation (Ostmeyer, Chakrapani, Pan, Perozo, & Roux, 2013). We reasoned that while D_2_O may affect channel gating via a non-specific mechanism – resulting from its increased viscosity (Kestin et al., 1985) or its effect on gating transitions (Fig. S6E, see also Díaz-Franulic et al., 2018), since recovery from slow inactivation takes place at resting membrane potential, at which minimal gating transitions takes place, it is unlikely to be affected by these non-specific effects. Comparison between the rates of recovery from slow inactivation of the M448K mutant in D_2_O and H_2_O based solutions revealed a small (18%) yet highly significant reduction in the fast component upon transition from H_2_O to D_2_O (Fig. S6D), consistent with a slower rate of exchange of D_2_O molecules buried behind the selectivity filter. Cs1 block resettling curves revealed a clear decrease in rate upon transition from H_2_O to D_2_O (Fig. 6B). This effect was specific, as similar experiments conducted with the classic pore-blocker ShK did not reveal any effect of D_2_O on block onset (Fig. 6C). The specific effect of H_2_O to D_2_O exchange on the development of block by Cs1 strongly suggest the existence of a step involving the motion of solvent molecules, independent from toxin binding, in the action mechanism of Cs1, as predicted from MD simulations.

**Figure 6:**
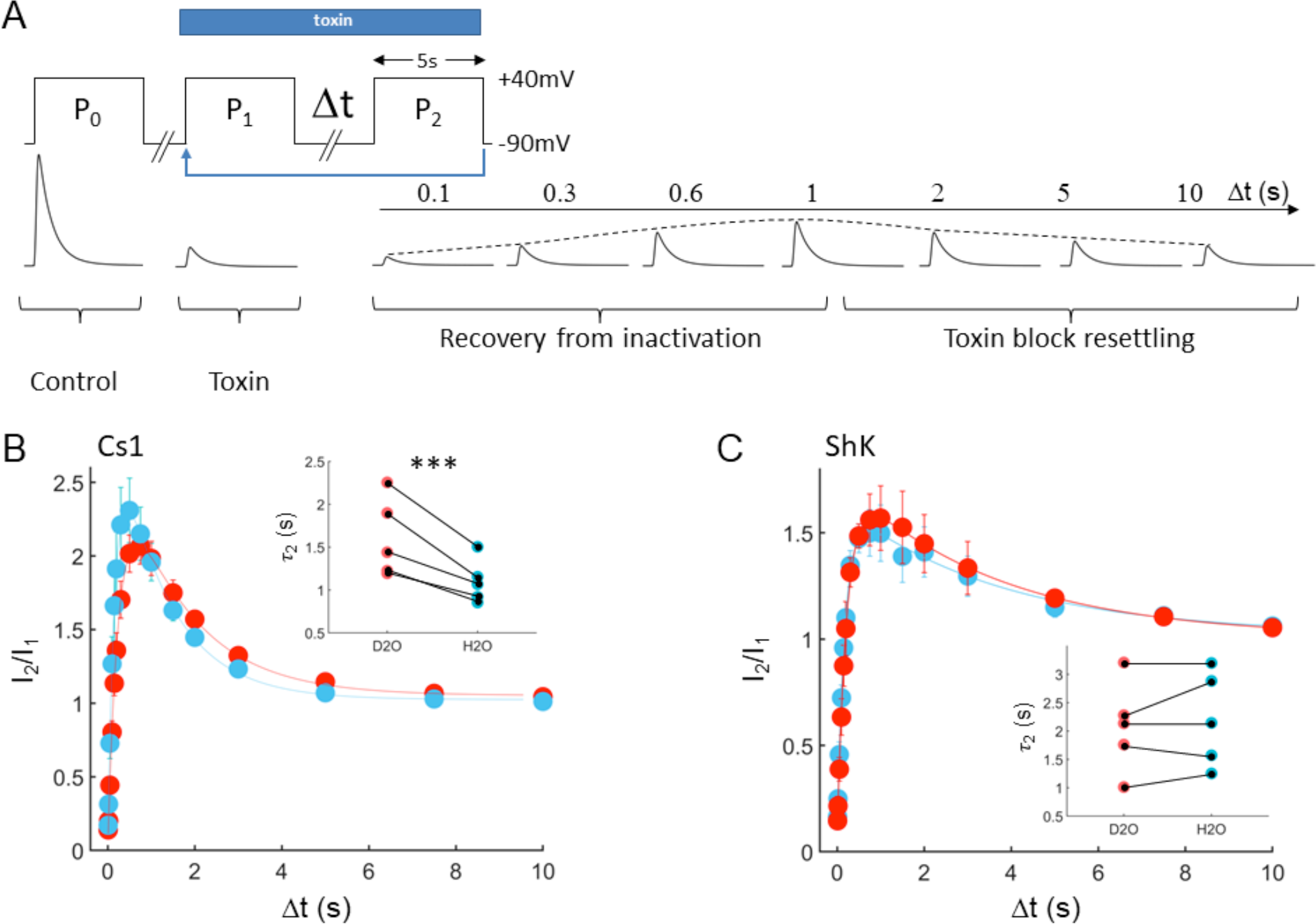
Channel block by Conkunitzin-S1 is slowed by heavy water. **A**, Experimental design allowing determination of channel block kinetics with high temporal resolution. The voltage protocol is presented at the top, current traces illustrating a typical response are presented at the bottom. A control pulse (P_0_) is applied to oocytes expressing Shaker_KD_ M448K, followed by application of a toxin dose inducing 80-90% block, and a first test pulse to +40mV (P1). During P1 channels enters a slow-inactivated non-conductive state and partial dissociation of the bound toxin takes place (Fig. S6 C,D). A subsequent recovery period of variable duration (Δt) at holding membrane potential (−90mV) allow channels to become available for re-opening (*τ* ~ 0.2s, Fig. S6B), but also promote re-binding and block by the toxin (*τ* ~ 2s). These two opposing processes affect the current amplitude measured at P2, which exhibit a bell-shape kinetics (dashed line), with a decaying phase that reports the rate of block onset (*τ*2, Eqn. 4). **B, C**, Channel re-block kinetics measured in the presence of 20nM ShK (right) or 50nM Cs1 (left). I_1_,I_2_ are the current amplitudes recorded during P_1_ and P_2_, respectively. Data were collected in either H_2_O (blue) or D_2_O (red) based solutions. Solid lines are best fits to eqn. 4, the derived time constants are summarized in table S6. The insets display *τ*_2_ values obtained from individual cells in H_2_O (blue) or D_2_O (red), solid lines connect data points derived from the same cell.

## Discussion

Animal toxins acting on voltage gated ion channels were forged by evolution to rapidly alter their targets, endowing venomous organisms with powerful tools for predation and defense. These toxins have traditionally been categorized into two classes – “gating modifier” toxins, which alter ion conduction by interfering with the voltage dependent motions of the channel, and “pore blocker” toxins that bind the channel pore and physically occlude the ion permeation pathway. Here we present a novel mechanism utilized by animal toxins to block K^+^ channels, which defies the traditional classification and expose a new pharmacological target present in most ion channels.

### Pore-modulation - a novel block mechanism of K^+^ channels

Classical pore blockers isolated from venomous organisms have played pivotal roles since the early days of ion channel research, facilitating the initial purification of novel channel proteins and providing the first insights into their subunit arrangement and the overall topology of the channel pore (Mackinnon, 1991; MacKinnon and Miller, 1989; Ranganathan et al., 1996). Canonic K^+^ pore blockers display high phylogenetic diversity and variable backbone folds; yet, they all share a strictly conserved lysine residue that physically plugs the channel pore (Dauplais et al., 1997). This mode of action was initially deduced from classical biophysical experiments (Park and Miller, 1992) and later gained strong support upon the elucidation of a solid-state NMR (Lange et al., 2006) and crystal (Banerjee et al., 2013) structures of channel-toxin complexes. Alongside the canonic K^+^ blockers, exists a diverse group of animal toxins that binds at the turret regions of K^+^ channels and blocks the ion conductance without physically occluding the pore (Hu et al., 2014; Rodríguez De La Vega et al., 2003; Verdier et al., 2005; Xu et al., 2003; Zhang et al., 2003). The precise molecular mechanisms by which toxins of this family block their targets have remained thus far elusive. Cs1, described herein, clearly belongs to this second group of K^+^ blockers and its novel mode of action, which we term “pore-modulation”, sheds a new light on the block mechanisms exploited by these toxins.

MD simulations of Shaker_KD_ in the presence of bound Cs1 revealed that in lack of tight binding between the toxin and the channel pore, water molecules and hydrated ions can access the extracellular vestibule (Fig. 2A). The demonstration that the non-inactivating Shaker_KD_ T449Y mutant exhibit normal ion conduction while bound to Cs1 (Fig. 5) further negates a “pore-lid” mechanism for this toxin. Instead, we suggest a mechanism in which the contacts between Cs1 and the channel turrets serve primarily to coordinate its critical interactions with the D-W gates. These gates consist of a network of hydrogen bonds that govern water permeation into the water-filled peripheral cavities that surround the central ion pore. Toxin interference with the D-W gates creates imbalanced water traffic at the peripheral cavities and triggers pore collapse. These three principal properties of Cs1 – turret binding, requirement for intact slow-inactivation and interaction with the D-W gates are shared by multiple K^+^ blocker families, for which a clear block mechanism has not been described. First. the cone-snail toxins of the κM family that exhibit many pharmacological similarities to Cs1, despite their radically different backbone fold. κM-conotoxin RIIIK of this group binds the turret region of the Shaker channel, does not block the non-inactivating T449Y mutant (Ferber et al., 2003), and exhibit a ring of positively charged residues directed towards the channel pore (Verdier et al., 2005). Second, the HERG blockers of the γ-KTX family. These scorpion toxins form multiple interactions with the extensive turrets of the HERG channel, yet, their block is strongly affected by substitutions at the upper-barrier that inhibit channel inactivation (Hu et al., 2014; Rodríguez De La Vega et al., 2003; Zhang et al., 2003). The recent structure of one such channel mutant, HERG:S631A, clearly demonstrate that the substitution prevents a tilt of the aromatic ring of the signature motif (G**F**G) and stabilize the canonic pore conformation (Wang and MacKinnon, 2017). Our analysis argues strongly that these toxins, alongside other “turret-binding” toxins, block the channel using the water-mediated pore-modulation mechanism described herein. Combined with recent reports of small molecules that bind at the peripheral cavities and modulate channel conductance (Lolicato et al., 2017; Wang and MacKinnon, 2017), pore-modulating toxins highlight these channel regions as an attractive pharmacological target and provides necessary cues for the rational design of compounds directed at these sites.

### Pore-modulation and slow inactivation

MD simulations of Cs1-bound Shaker_KD_ reveal a sequence of events that culminate in an asymmetric collapse of the channel pore, triggered by the bound toxin. Several recent studies conducted with unmodified and chemically modified channels suggest that the molecular events captured by our simulations faithfully describe channel dynamics. First, chemical modifications at the SF of KcsA that enhance slow inactivation were shown to trigger asymmetric collapse of the channel pore, by altering the turnover of water molecules bound behind the SF. Basins at the free energy landscape of unmodified KcsA, corresponding to the asymmetrically constricted pore conformation were detected (Li et al., 2017). Second, a chemical modification of K_v_1.2:Trp434 was shown to enhance slow inactivation by increasing water traffic through the D-W gate (Lueck et al., 2016). A key event that re-occurred in all Cs1-bound simulations was the collapse of the hydrophobic cuff barrier that enabled water exchange between the peripheral pore and the central vestibule of the channel and between pockets of neighboring subunits (Fig. S4 F,H, movies S4B,C). Flips of the pore backbone carbonyls then diverted this water stream into the main pore, leading to ion loss and rapid collapse of the SF. Flipped backbone carbonyls were captured in the crystal structures of inactivated ion channels (Cordero-Morales et al., 2006). The recent Cryo-EM structures of the HERG channel and its non-inactivating S631A mutant further establish a direct link between a compromised cuff-barrier and the inactivated state of the channel (Wang and MacKinnon, 2017). Water conduction behind the collapsed SF of KcsA was suggested to explain the high water permeation of its closed state (Furini et al., 2009; Hoomann and Jahnke, 2013). Our simulations reveal an opening of a hydrophilic path connecting the peripheral cavity with the intracellular vestibule that involves two conserved threonine sidechains (Fig. S4F). Alanine substitutions at the second of these Thr residues were recently shown to impair slow inactivation in K_v_1.2, K_v_1.5 and KcsA and to alter the occupancy of water molecules buried behind the SF of crystalized KcsA mutants (Labro et al., 2018). Combined, these studies portray a molecular machinery composed of water paths and gates behind the K^+^ channel pore that set the rate and timing for slow inactivation. Pore modulating toxins efficiently exploit this machinery to impose ectopic channel gating.

## Materials and Methods

### Animals

Female wild-caught *Xenopus leavis* frogs were purchased from NASCO. Experiment were performed according to the guidelines of the Weizmann Institute Animal Care and Use Committee permit #37070717-3.

### Method details

#### Expression and purification of Conkunitzin-S1 and ShK

E. coli BL21 (DE3) were transformed with the appropriate toxin construct cloned into the SapI/BamHI sites of pTwin1 (New England Biolabs). Freshly transformed colonies were used to inoculate a 10-ml starter in LB containing 0.1mg/ml Carbenicillin and 0.03mg/ml chloramphenicol. Following 12 hours of growth at 37°C, the starter was diluted into a baffled flask containing 1 liter LB supplemented with antibiotics and grown to OD_600_ of 0.6. IPTG was added to a concentration of 0.4 mM and culture growth was continued for additional 5 hours. Cells were harvested by centrifugation (10 min at 5000 x g at 4°C) and re-suspended in 50 ml H_2_O. The cell suspension was frozen and thawed at 50°C for 45 minutes to disrupt cell walls and total lysis was achieved by sonication. The cell lysate was centrifuged for 15 min at 20,000 x g and the inclusion bodies pellet was washed and precipitated twice in washing solution containing 25% (W/V) sucrose, 5 mM EDTA, 1% (V/V) Triton X-100, in PBS pH 8.0. The inclusion bodies were dissolved and incubated for 1 hour in 10 ml denaturation buffer containing 6 M Guanidinium-HCl, 0.1 M Tris-HCl pH 8.0, 1 mM EDTA and 10mM reduced glutathione. The denatured protein suspension was diluted 20-fold into cold (4°C) renaturation solution composed of 0.2 M ammonium acetate pH 8.0, 0.5M Sucrose and 0.2 mM oxidized glutathione, and incubated at 4°C with gentle stirring for 24 hours. The renaturation mixture was filtered through 1 mm Whatman paper to eliminate insoluble proteins and the soluble material was precipitated in 50% ammonium sulfate at 4°C for 4 hours. The precipitate was collected by filtration through a GF/C membrane and re-suspended in 20ml phosphate buffer (50mM, pH 6.2) to activate intein cleavage which releases the toxin from the tag. Following 24 hours incubation at room temperature, the soluble protein was isolated by centrifugation and 0.2μM filtration. The protein sample was then subjected to RP-HPLC, using a 10ml tricorn column packed with source 15RPC on an ACTA-Pure HPLC system. Proteins were eluted with a linear gradient of acetonitrile containing 0.1% TFA (buffer B). Cs1 typically eluted at 28% B and ShK at 23% B, enabling clear separation from the intein moiety that eluted at 70% B. RP-purified samples were dried by lyophilization and kept at −80°C.

#### Toxin crystallization, X-ray data collection and structure determination

Crystals of Cs1 Q54A and Cs1 R55D were grown using the hanging-drop vapor-diffusion method. The crystals of Cs1 Q54A were grown from 0.2 M Ammonium sulfate, 0.1 M Sodium acetate trihydrate pH 4.6 and 25% w/v PEG 4,000. The crystals formed in the hexagonal space group *P63*, with one molecule per asymmetric unit. Crystals of Cs1 R55D were grown from 0.05M Potassium phosphate monobasic and 20% w/v PEG 8,000. The crystals formed in the trigonal space group *P3_2_21*, with two copies per asymmetric unit. Complete datasets to 1.3Å resolution of both protein crystals were collected at 100K using ADSC Q4 CCD detector at ID14-EH4 of the European Synchrotron Radiation Facility (ESRF, Grenoble, France).

Molecular replacement: The structures of the toxins were solved by molecular replacement using the program MOLREP (Vagin and Teplyakov, 2000). The search model was based on a low resolution structure of Conkunitzin-S1 previously determined at 2.4 Å resolution (PDB code 1Y62). All steps of atomic refinement of both structures were carried out with the CCP4/REFMAC5 program (Murshudov et al., 1997) and by Phenix refine (Afonine et al., 2012). The models were built into *2mF*_*obs*_ - *DF*_*calc*_, and *mF*_*obs*_ - *DF*_*calc*_ *maps* by using the COOT program (Emsley and Cowtan, 2004). Details of the refinement statistics of the Cs1 Q54A and Cs1 R55D structures are described in **Table 1**. The coordinates of Cs1 Q54A and Cs1 R55D were deposited in the RCSB Protein Data Bank with accession codes 6Q61 and 6Q6C respectively. The structures will be released upon publication.

**Table 1:** X-ray Data Collection Statistics for the Cs1 54A and Cs1 55D crystals.

|  | **54A** | **55D** |
| --- | --- | --- |
| **PDB Code** | 6Q61 | 6Q6C |
| **Data collection** |  |  |
| Space group | *P*63 | *P* 3<sub>2</sub> 2 1 |
| Cell dimensions: a,b,c (Å) | 51.03, 51.03, 42.57 | 64.83, 64.83, 66.82 |
| Resolution (Å) | 19.61-1.30 | 28.07-1.3 |
| Upper resolution shell (Å) | 1.347-1.301 | 1.346-1.3 |
| Measured reflections<sup>a</sup> | 280,741 | 148,645 |
| Unique reflections | 15,585 (1,554) | 40,174 (3,932) |
| Multiplicity | 10.8 (10.7) | 3.7 (3.7) |
| Completeness (%) | 99.78 (99.61) | 99.41 (98.84) |
| Average I/ $\sigma$ (I) | 24.99 (2.40) | 19.75 (2.79) |
| Wilson B-factor (Å<sup>2</sup>) | 15.62 | 16.29 |
| R-pim | 0.01299 (0.2929) | 0.06148 (0.714) |
| **Refinement Statistics** |  |  |
| Reflections used in refinement | 15,565 (1,551) | 40,150 (3,932) |
| Reflections used for R-free | 779 (76) | 2,019 (192) |
| R-work | 0.1490 (0.1996) | 0.1673 (0.2964) |
| R-free | 0.1788 (0.2237) | 0.2034 (0.3110) |
| Number of non-hydrogen atoms | 566 | 1162 |
| Number of protein atoms | 509 | 977 |
| Solvent | 52 | 147 |
| RMS(bonds) (Å) | 0.017 | 0.055 |
| RMS(angles) (°) | 1.97 | 2.24 |
| Ramachandran favored (%) | 96.49 | 97.35 |
| Ramachandran allowed (%) | 3.51 | 2.65 |
| Ramachandran outliers (%) | 0.0 | 0.94 |
| Average B-factor (Å<sup>2</sup>) | 29.34 | 26.11 |

<sup>a</sup>The values in parentheses refer to the data of the corresponding upper resolution shell

#### Circular dichroism spectrometry

CD-spectra were recorded at 25°C using a model 202 circular dichroism spectrometer (Aviv Instruments, Lakewood, NJ, USA). Toxins (140 μM) were dissolved in 5 mM sodium phosphate buffer, pH 7.0, and their spectrum measured using a quartz cell of 0.1 mm light path. Blank spectrum of the buffer was run under identical conditions and subtracted from each of the toxin spectra.

#### Electrophysiology – oocytes handling

Plasmids encoding for the Drosophila Shaker and its mutants clone in pBlueScript were linearized using *EcoRI* and purified using phenol/chloroform. The linearized templates were used for mRNA synthesis using the T7 mMESSAGE mMACHINE Transcription Kit (Ambion) and stored as stock solutions at −80°C. *Xenopus leavis* female frogs were anesthetized and subjected to surgery by incision in the lower half of the belly. The oocytes were pulled out from the incision and placed in sterile calcium free ND96 solution containing 96mM NaCl, 2mM KCl, 1mM MgCl_2_ in 5mM HEPES pH 7.5 and then incubated in collagenase solution (3mg/ml) for 2 hours in order to defoliculate them. After the collagenase treatment, the oocytes were incubated in ND96 solution supplemented with 1.8 mM CaCl_2_, 2.5 mM sodium pyruvate, and 100 mg/ml gentamicine (NDE). Selected defolliculated oocytes were injected with 1ng mRNA of interest using a Drummond 510 microdispenser. Injected oocytes were incubated at 18°C for 1-2 days in NDE solution prior the electrophysiological experiments.

#### Electrophysiology – data acquisition

K^+^ currents from the injected *Xenopus* oocytes were measured by two-electrode voltage clamp using a Gene Clamp 500 amplifier (Axon Instruments, Union City, CA, USA). Oocytes were placed in a 100 μl fiberglass bath and perfused with ND96. Toxin solutions were freshly made by the re-suspension of lyophilized toxin in ND96 supplemented with 1 mg/ml BSA, and then diluted and applied directly to the bath. Some measurement that required high external K^+^ were conducted in a 90K solution containing 90mM KCl, 2mM MgCl_2_, 10mM HEPES at pH 7.4. Experiments in which the effects of D_2_O were examined were conducted in ND96 or 90K solutions freshly prepared using D_2_O as the solvent. Data were sampled at 10 kHz and filtered at 5 kHz using a Digidata 1550A device controlled by pCLAMP 10.5 (Axon Instruments, Union City, CA). Capacitance transients and leak currents were removed by subtracting a scaled control trace utilizing a P/4 protocol.

#### Electrophysiology – Data analysis

**Dose-response curves** were acquired by the application of increasing toxin concentrations into the measurement bath. Each curve was constructed from at least 5 toxin concentrations, an adequate period of incubation (> 1 min) at each toxin concentration ensured steady-state response. Experiments were carried in triplicates. Data points were fit using a Hill equation with the hill coefficient restricted to unity:

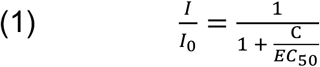

Were I_0_ is the unmodified current measured before toxin application, C is toxin concentration and EC_50_ is the half maximal toxin concentration. Dose-response curves of Cs1 were acquired on the background of the K427D mutation, unless otherwise noted, and this channel mutant is referred throughout the manuscript as Shaker_KD_. The high affinity binding of the toxin to this channel allowed for exact potency determinations by achieving effect saturation and facilitated double-mutant cycle analysis using channel mutants with low affinity for the toxin.

**Double-mutant cycle analysis** – Toxin residues that were found important for toxicity (R34, R49, Y51, R55 and Y59) were substituted to alanine, and the potencies of the resulting mutants were assayed on each of the high impact channel mutants (F425A, D427K, D431A). Coupling energies between toxin and channel residues were extracted from the experimental data as described by Hidalgo and MacKinnon (1995), assuming a linear relation between toxin binding and channel block (ie, EC_50_ = K_d_). For each pair of channel/toxin mutants a coupling coefficient was calculated:

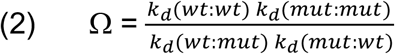

from which a coupling energy was extracted by

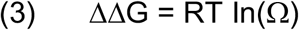

where R is the gas constant, and T is the temperature (25°C).

**Voltage-dependent toxin dissociation assays** - Oocytes expressing Shaker_KD_ were exposed to 2nM ShK or 5nM Cs1 for 5 minutes until a stable block of ~80% was achieved. Voltage dependent toxin-dissociation was assayed using a two-pulse voltage protocol (Fig. 2C, inset) in which two test pulses (P1,P2) of 50ms to −10mV applied from a holding potential of −90mV were separated by a strong depolarizing pulse (+100mV) of variable duration. Voltage-dependent dissociation of the bound toxin leads to an elevated amplitude of the current recorded during the second test pulse and an increase of the ratio P2/P1.

**Channel re-block assays** – oocytes expressing Shaker_KD_ M448K with current densities of 5-7μA were subjected to toxin dose inducing 80-90% block and the toxin effect was allowed to settle for at least 1 minute. Bell-shaped curves of current amplitude during channel recovery from inactivation and re-block by the toxin were acquired as described in fig. 6. The data were fit by a bi-exponential function:

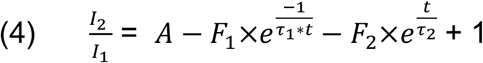

Where I_1_ and I_2_ are the current amplitudes at P_1_ and P_2_, respectively, and τ_1_ and τ_2_ are the rate constants for recovery from inactivation and re-block by the toxin, respectively.

### MD simulations

#### Modeling of the Shaker_KD_-Cs1 and Shaker_KD_-ShK complexes

A structural model of Shaker_KD_ pore domain (residues Lys376-Asp488) was constructed based on the published crystal structure of the K_v_1.2-K_v_2.1 paddle chimera (PDB ID 2R9R, 86% identity, 94% similarity) using MODELER (Webb and Sali, 2017). For docking Cs1, the 1.3Å crystal structure of the highly active (EC_50_ = 3.57±0.74nM, n=3) Q54A mutant (table 1, PDB ID 6Q61) was utilized. For the docking of ShK, we have utilized the published solution structure of the toxin (Tudor et al., 1996). The resulting models were submitted for a rigid-body docking without any constraints using the CLUSPRO web service (Kozakov et al., 2017), and the solutions were ranked using the weighting coefficients of the Van der Waals+electrostatics docking mode. For both toxins, the majority of the top 1000 solutions were divided between four clusters, which could be converged into a single unique solution by rotation about the pore axis. These unique solutions were further validated by cross-referencing against the available experimental restraints (for Cs1, see Results, for ShK see Lanigan et al., 2002; Rashid and Kuyucak, 2012). For modeling the interaction of CTX with the K_V_1.2-K_v_2.1 paddle chimera, we utilized its available crystal structure (4JTA). The simulation system for the Shaker_KD_ T449Y:Cs1 was constructed by the sequential substitution of T449 in all four subunits of the pre-assembled Shaker_KD_:Cs1 system using the MUTATOR plugin of VMD.

#### MD System setup

The simulation systems employed in this study were assembled using the VMD software suite (Humphrey et al., 1996). The channel protein was embedded in a lipid bilayer containing 190 POPC molecules and solvated with 11621 (toxin free system) to 13773 (toxin bound systems) water molecules. Two K^+^ ions were initially placed at sites s2 and s4 and a water molecule was placed at site s3 of the selectivity filter based on the crystallographic coordinates to obtain an initial 2,4 configuration. Additional K^+^ and Cl^−^ ions were added to neutralize the system and set the KCl concentration at 150mM. Final system dimensions were roughly 95 X 95 X 80 Å^2^ for toxin free and 95 X 95 X 100 Å^2^ for toxin bound systems. We have used the CHARMM36 parameter set for proteins, lipids and salt ions, and the CHARMM TIP3P model for water (Best et al., 2012).

#### MD Simulation protocol

Simulations were carried either on GPU using the Win64/CUDA, or on CPU using the Linux-x86_64-ibverbs versions of NAMD (Phillips et al., 2005). All simulations were performed under NPT conditions at 300K and 1atm. Periodic boundary conditions and electrostatic interactions were treated using the particle mesh Ewald method with ~1Å grid size and a 12Å cutoff distance. A time step of 2fs was employed, and bonds involving hydrogen atoms were fixed using the SHAKE algorithm. Assembled systems were minimized using 2000 steps of a conjugated gradient and line search algorithm and then equilibrated as follows: initially the protein coordinates were fixed and the system was equilibrated for 0.5ns to allow the lipids and the solvent to assume proper densities. Then, a series of 0.2ns equilibration steps with decreasing harmonic restrains of 10, 5, 2, 1, 0.6, 0.3 and 0.1 Kcal/mol/Å^2^ was applied first to the sidechains, and then to the backbone of the protein. System equilibration was concluded by an unconstrained 20ns run. Six unique systems were simulated in the scope of the current study. Each production run was 200ns long. Seven different production runs were carried for the Cs1: Shaker_KD_ complex; the other systems, namely, Cs1: Shaker_KD_ T449Y, ShK: Shaker_KD_, CTX: Paddle Chimera, free Shaker_KD_ and free Shaker_KD_ T449Y, were each simulated in 3 independent production runs.

#### Analysis protocols for MD simulations

Trajectory snapshots were saved every 100ps during production simulations. Custom written MATLAB code was used for trajectory analysis (see supplementary file). The peripheral cavity of a given channel subunit was empirically defined as the volume formed by the intersection of four spheres, with centers coordinates at the backbone nitrogen atoms of Val438, Tyr445, Met448 and Val451 and with radii of 10, 7, 10 and 14Å respectively. This definition effectively filtered out pore-waters and water molecules from the neighboring cavities. For RMSF calculations, the mean position of each water molecule during its residence period in a given cavity was used as the reference distance. The radius of gyration for a backbone carbonyl of a given residue was calculated as the distance of its oxygen atom from the geometrical center of the C_α_ atoms contributed by that residue in each of the four subunits. Hydrogen bonds were scored based on a donor-acceptor distance of less than 3Å and a cutoff angle of 120°.

#### Quantification and Statistical Analysis

All statistical analyses were conducted using the MATLAB built-in functions ttest or ttest2 for one sample/paired sample T-tests. All data are expressed as the mean ± standard deviation, unless otherwise stated in the figure legend.

## Supplementary figures

**Figure s1:**
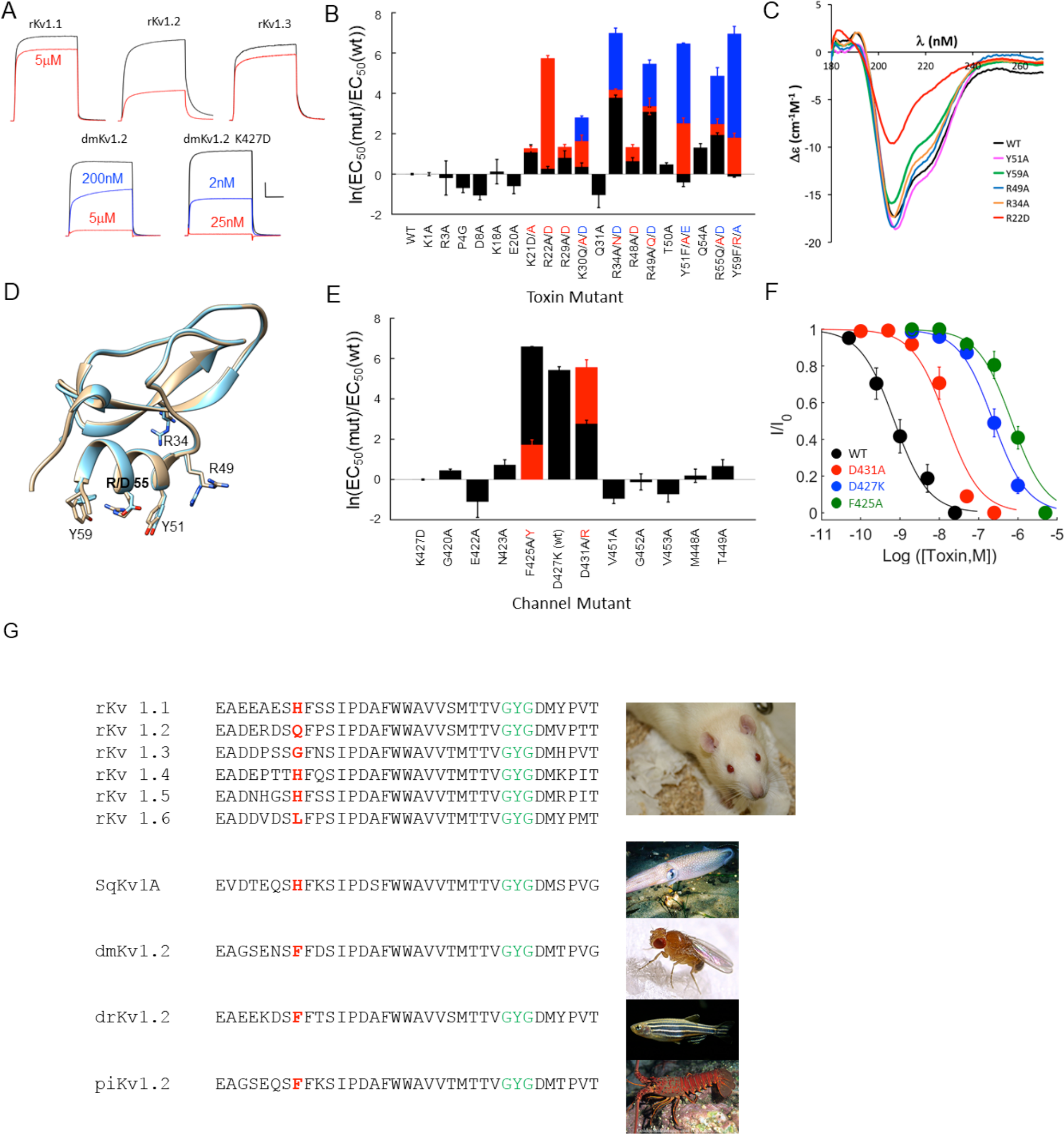
molecular and functional dissection of the Shaker_KD_-Cs1 complex. **A**, The differential preference of Cs1 for mammal (top) and insect (bottom) K_v_s. The indicated channel subtypes were expressed in *xenopus leavis* oocytes, and the resulting currents were recorded using TEVC. Toxin effect was assayed by direct bath application. Traces in black were acquired before toxin application, colored traces were recorded in the presence of the indicated toxin concentrations. **B**, The impact of point mutations in Cs1 on its potency on Shaker_KD_. Data are reported as the natural log of the relative potency for each toxin mutant. In cases where multiple point mutations were introduced, different colors are used as indicated. **C**, CD spectra of high-impact Cs1 mutants. All mutants that were assigned to the bioactive surface of the toxin exhibit a wt-like spectra. The CD spectra of the R22D mutant is significantly altered, likely due to the abolishment of an intramolecular salt bridge. **D**, the crystal structure of Cs1 Q54A (tan, EC_50_ = 3.57±0.74nM) and R55D (cyan, EC_50_ = 132±50nM) are overlaid, bioactive residues are rendered in sticks. **E**, The impact of point mutations in Shaker_KD_ on its affinity for Cs1. **F**, Dose-response curves of Cs1 on the indicated Shaker_KD_ mutants expressed in oocytes. The EC_50_ values obtained were 0.95±0.03 nM (unmodified), 15.2±2.7nM (D431A), 219±37nM (D427K) and 693±18nM (F425A). G. The low affinity of Conk-S1 for mammalian K_v_s is dictated by the side chain at position 425. Sequence alignment of the pore-helix and membrane re-entrant loop region of six K_v_ channel isoforms from rat (rK_v_1.1-1.6), and representative clones from squid (SqK_v_1A, from *Doryteuthis opalescens*), insect (dmK_v_1.2 from *drosophila melanogaster*), fish (drK_v_1.2 from *Danio rario*) and a crustacean (piK_v_1.2 from *Panulirus interruptus*). Residues at position 425 are highlighted in red, the GYG signature sequence is highlighted in green.

**Movie S2:**
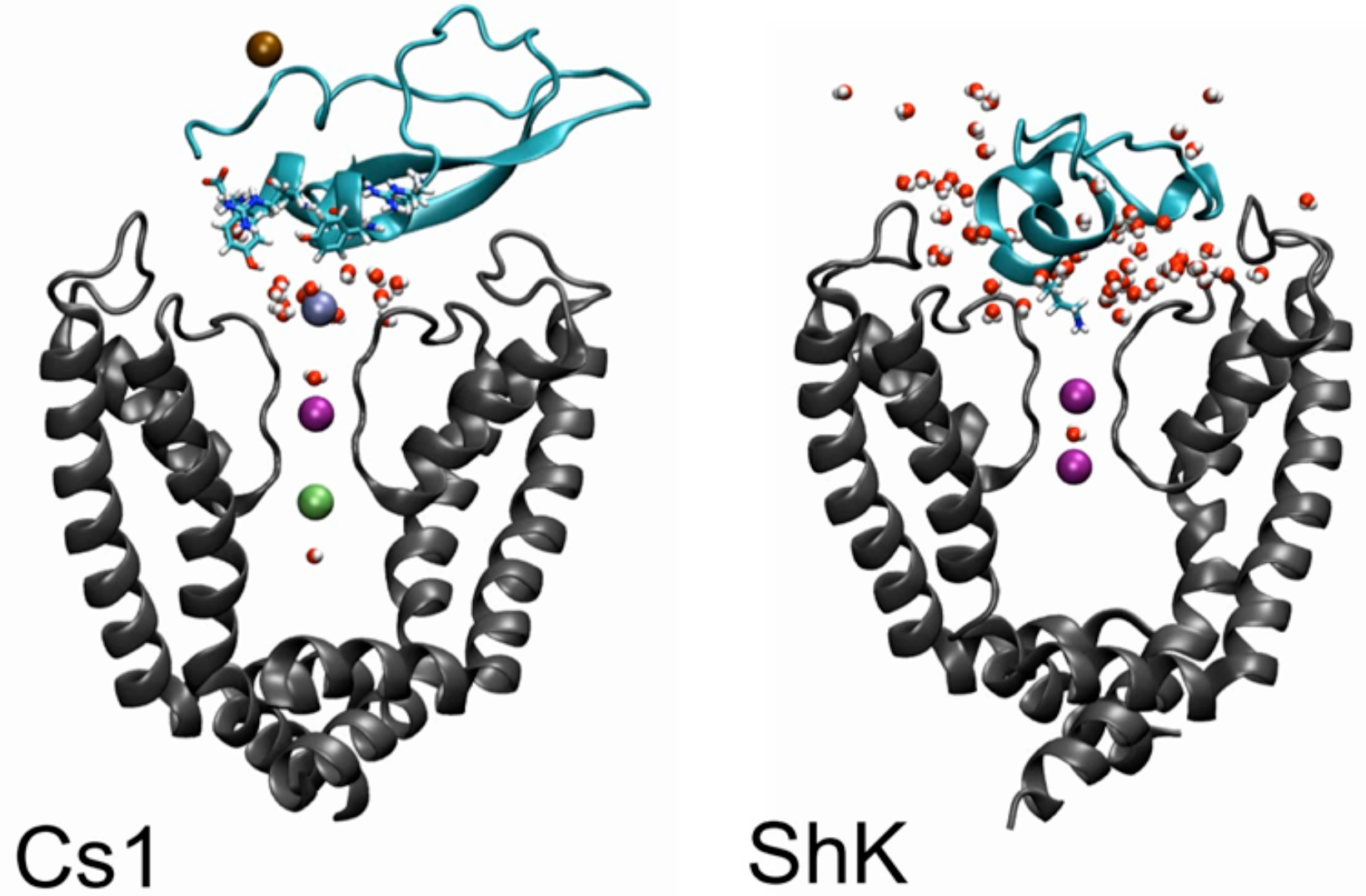
Conkunitzin-S1 does not directly block the ion conduction pathway. The movie depicts the traffic of water and K^+^ ions at the channel/toxin interface for the Cs1 and ShK toxins over a 50ns MD simulation window. The rendering convention is as in Fig. 2B. For the Cs1 trajectory, K^+^ ions are rendered in unique colors. For ShK, all water molecules within a cutoff distance of 5Å from both the toxin and the membrane plane are displayed. For Cs1, an additional criterion was applied – only water molecules that traversed the extracellular pore entry (Fig. 2B) during the simulation window are rendered.

**Figure S3:**
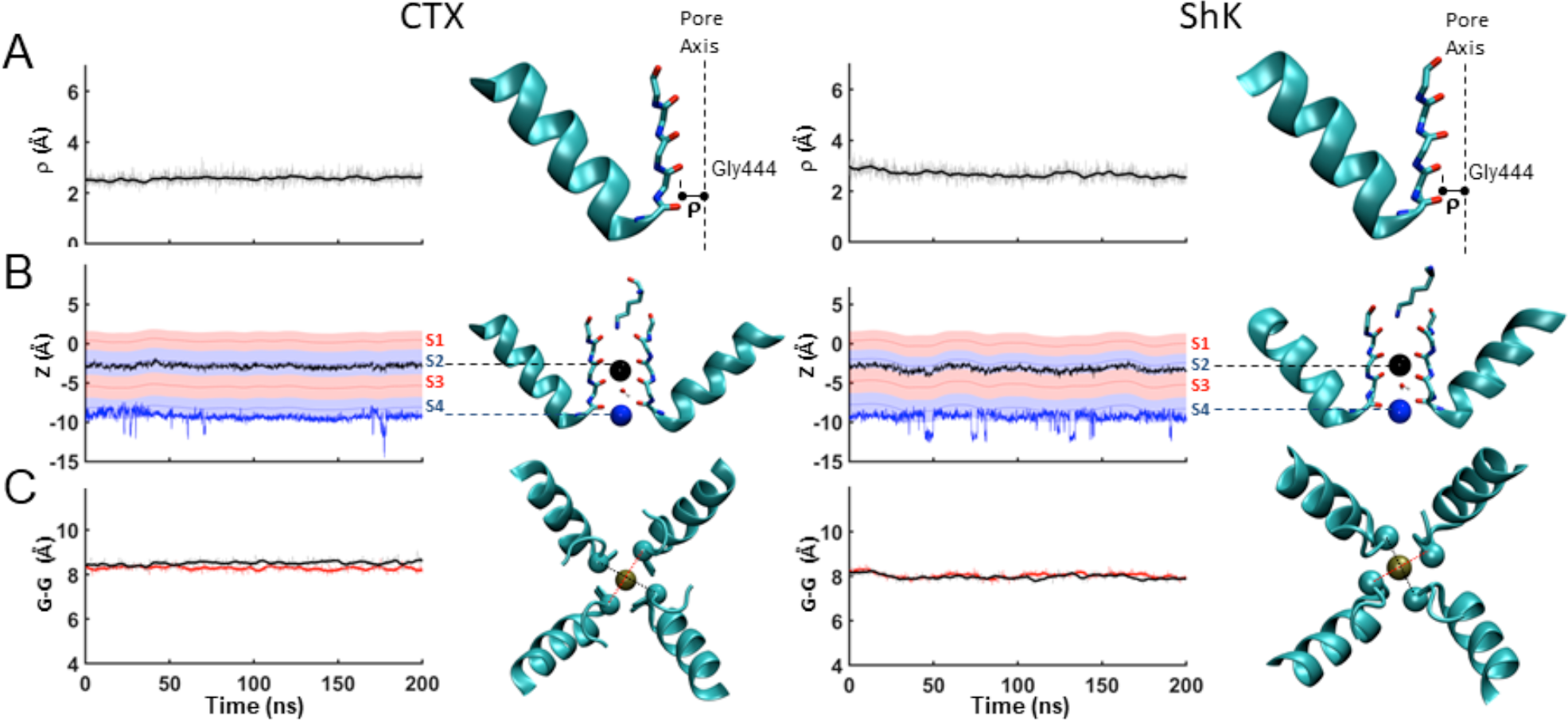
the conformation of the channel pore is maintained during simulations of bound classical K_v_ pore blockers. Conformational dynamics of the selectivity filter region during 200ns long trajectories of CTX bound to the paddle chimera (PDB ID: 4JTA, left) and ShK bound to Shaker_KD_ (right). For details see Figure 3.

**Figure S4:**
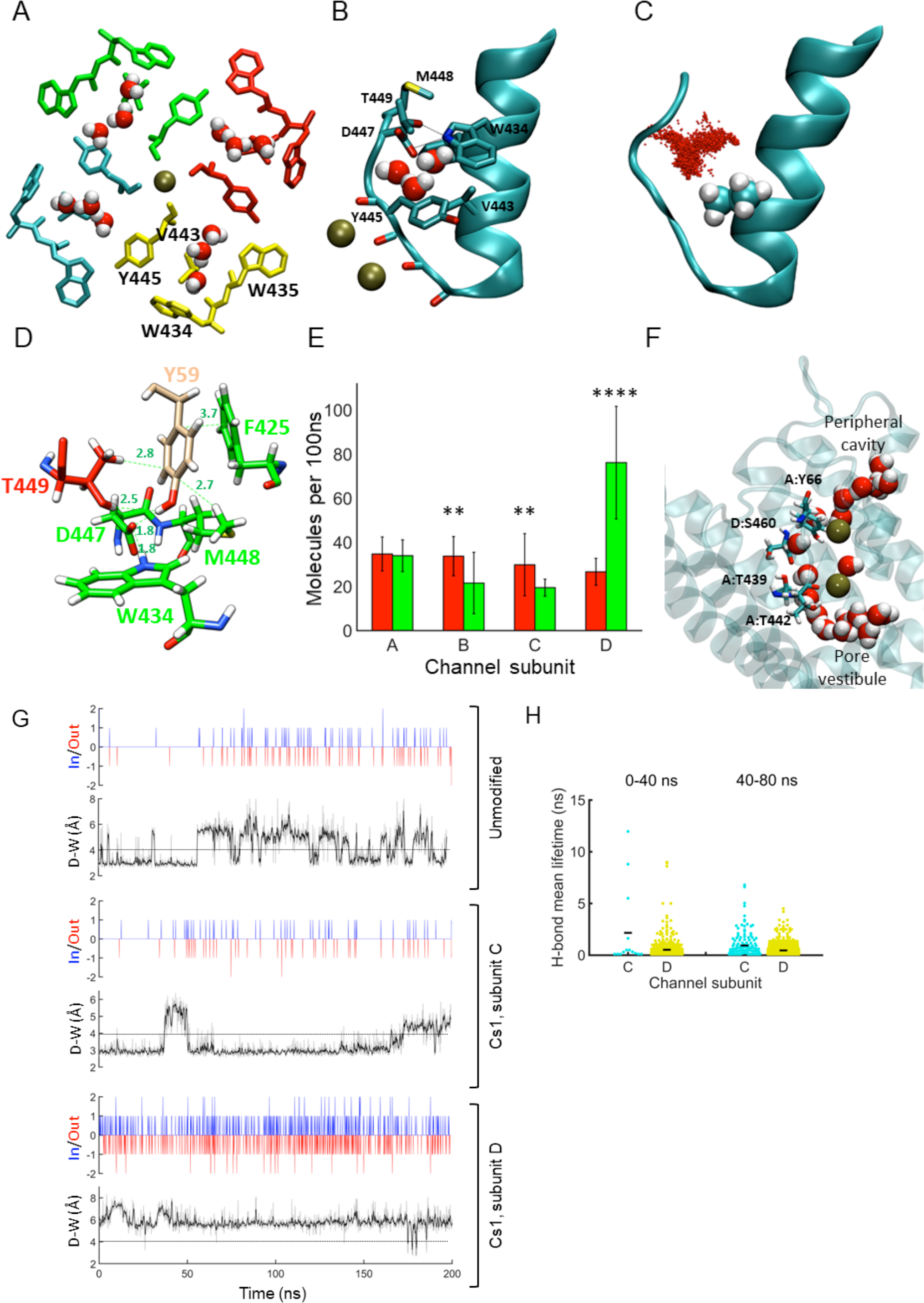
Conkunitzin-S1 modifies water permeation into the peripheral cavities. **A**, Top view of the aromatic-cuff barrier and the cavity water molecules of Shaker_KD_. A snapshot taken following a 200ns production run, depicting the sidechains of W434, W435, V443 and Y445 that constitute the lower barrier for water molecules movement inside the cavity. A by-chain coloring scheme apply (A-red; B-green; C-blue and D-yellow). Cavity water molecules and pore ions are rendered as spheres. **B**, Side view of a peripheral cavity, residues that constitute the upper and lower barriers for water movement are rendered in sticks. The bifurcated hydrogen bond between W434 N_ε_1 and D447-O_δ_1 / Thr449-O_γ_1 is indicated by dashed lines. **C**, a typical spatial distribution of water molecules within a peripheral cavity of unmodified Shaker_KD_ along a 200ns trajectory. Dots mark the centers of the water oxygen atoms. **D**, Snapshot taken at the end of 20ns equilibration run, depicting multiple interactions between Cs1:Tyr59 and the upper barrier residues in subunit B. Green labels indicate distances(Å). **E**, Cs1 increase the rate of water exchange at the peripheral cavities of chain D while decreasing it at chains B and C. The bars depict the number of unique water molecules passing through the peripheral cavities of unmodified (red) and Cs1-bound (green) channels during a 100ns simulation window. **F**, Hydrophilic pathway taken by cavity water molecules following the collapse of the aromatic cuff barrier (see movie S4B). The backbone of channel subunits A and D is rendered as a transparent ribbon. water molecules along a pathway that connects the peripheral cavity of subunit D through the breached barrier with the intracellular vestibule of the channel are rendered in spheres. Channel residues that coordinate these water molecules are rendered in sticks. **G**, The rate of water exchange between the peripheral cavities and the bulk solution is highly correlated with the D447-W434 gate state. Paired time-series plots depicting water exchange events (top) vs. D447:W434 gate openings (bottom) for unmodified channel (top), Cs1-bound channel subunit C (middle) and Cs1-bound channel subunit D (bottom) cavities are shown. In the water exchange plots, blue bars marks water entry events, while red bars mark water exit events. The DW-gate plots were prepared as described in Fig. 4B. **H**, Flip of the Y445 sidechain in domain D enables exchange of water molecules between the peripheral cavities of subunits C and D (related to movie S4B). Mean lifetimes of the hydrogen bonds formed by water molecules residing in the peripheral cavities of subunits C and D is used as a measure of mobility. Note the increased mobility at subunit C cavity following the ring flip that takes place at t = 40ns.

**Movie S4A:**
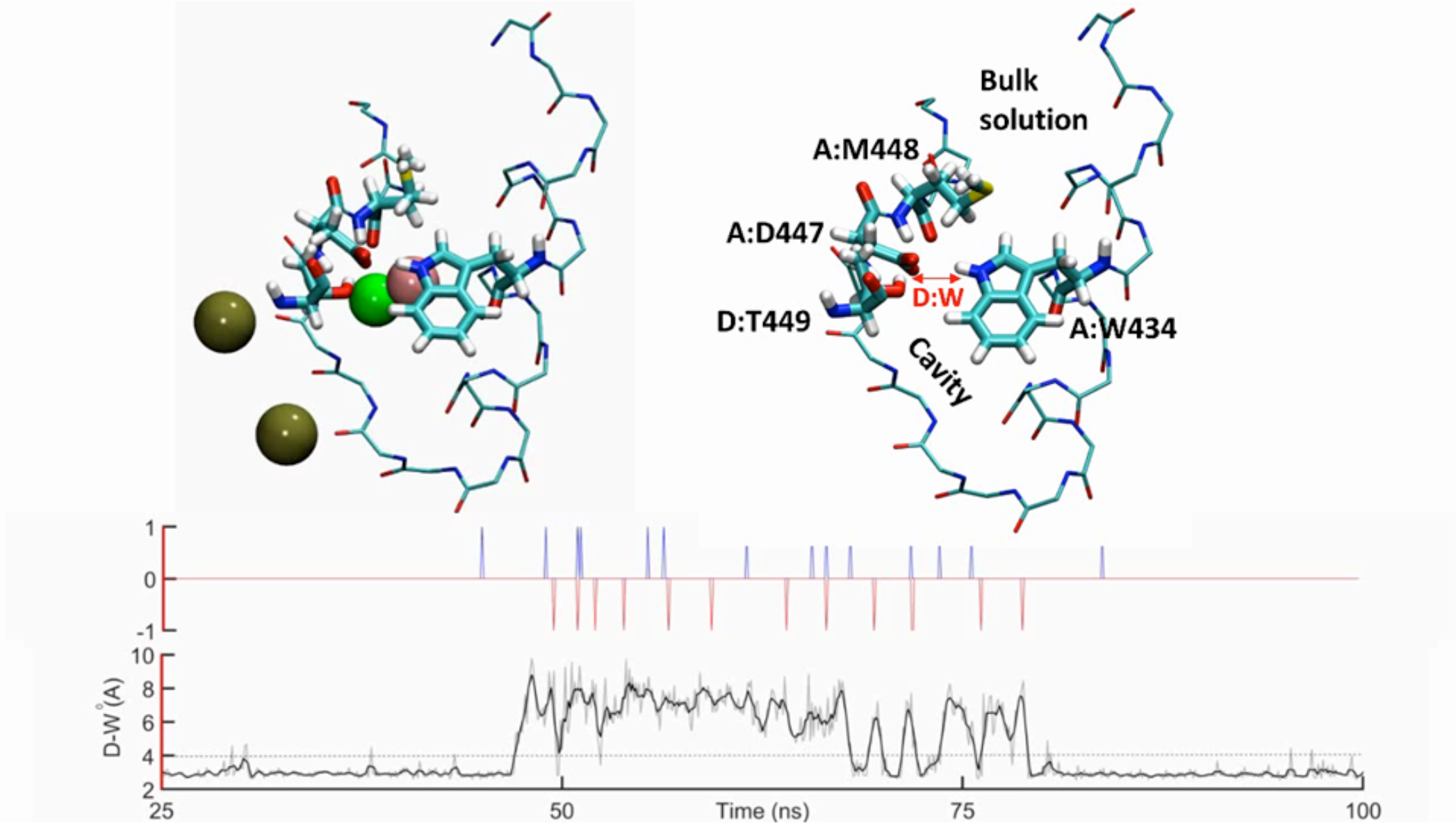
The rate of water exchange between the peripheral cavity and the bulk through the upper barrier is highly correlated with the D447-W434 gate state. The movie depicts the entry/exit of water molecules from a peripheral cavity of Shaker_KD_ during a 75ns simulation window. Backbone atoms of residues that line the cavity are rendered as thin sticks, residues of the upper barrier are rendered as thick sticks. Spheres depict pore ions (tan) and oxygen atoms of cavity water molecules (variable colors). Visual legend for the displayed residues and overall topology is provided at the right. Time series plots at the bottom depict water exchange events (top, blue for entry, red for exit), and the D-W gate state (The distance between the O_δ_1 atom of Asp447 and the N_ε_1 atom of Trp434). At t~45ns the DW gate switches from a closed to an open conformation, dramatically increasing the rate of water exchange.

**Movie S4B.**
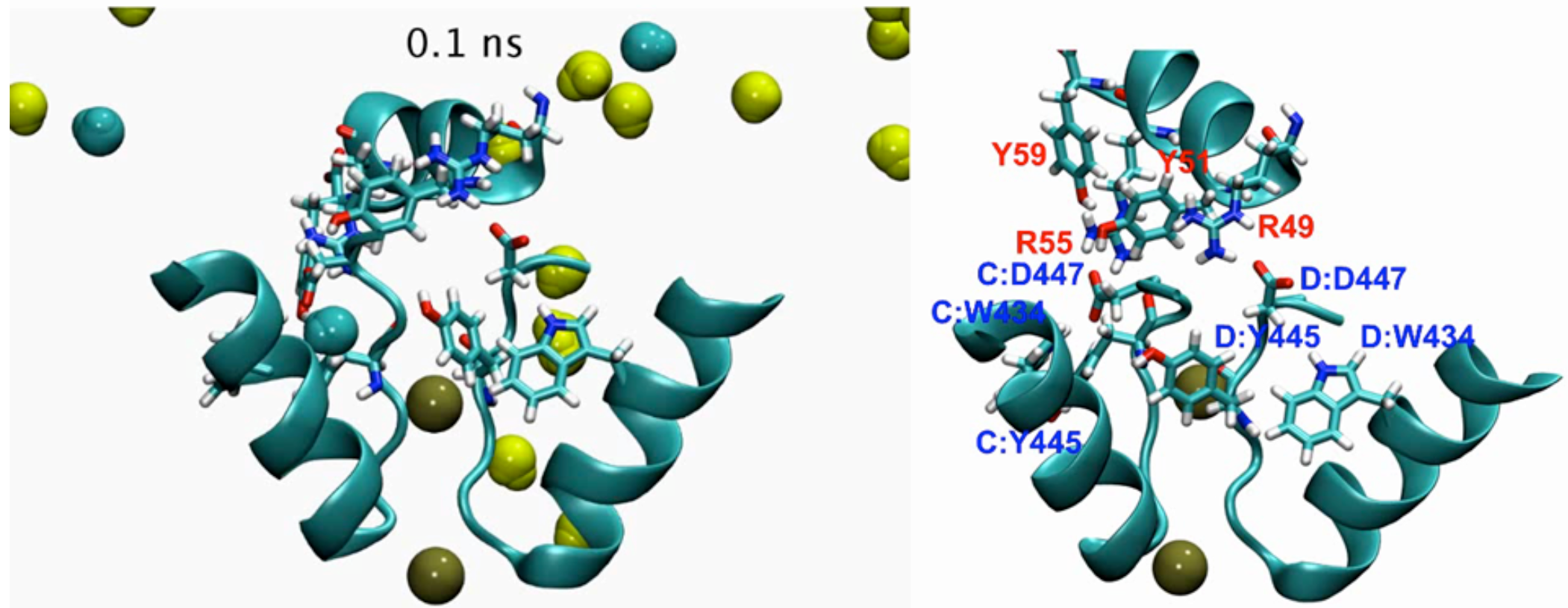
Flip of the Y445 sidechain in domain D enables exchange of water molecules between the peripheral cavities of subunits C and D. The pore helices of subunits D and A as well as the C-terminal α-helix of Cs1 are shown. The oxygen atoms of water molecules residing at the peripheral cavities D (yellow) and A (red), and K^+^ ions in the pore (tan) are rendered as spheres. Visual legend for the displayed residues is provided at the right. Note the flip of D:Y445 at t~40ns that enable water exchange with the peripheral cavity of chain C (Fig S4H).

**Movie S4C.**
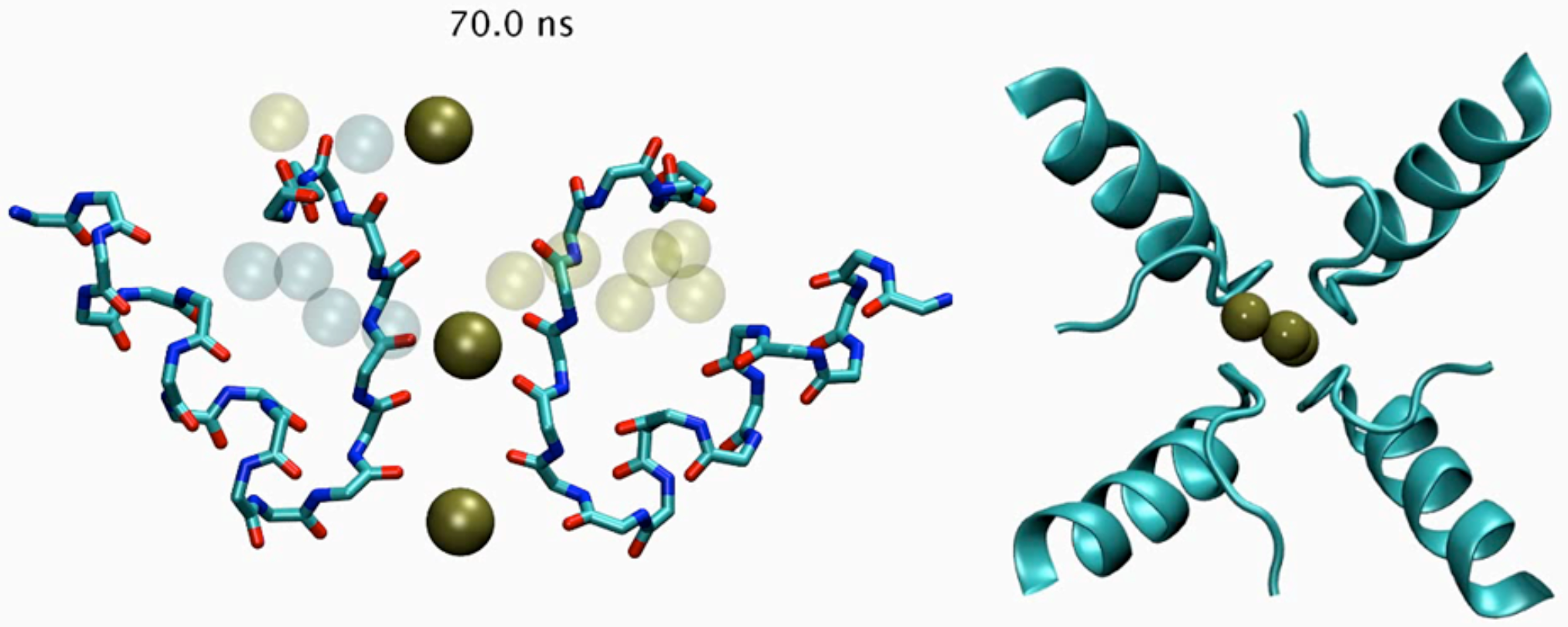
Invasion of water molecules from the peripheral cavities into the central pore displace bound K^+^ ions and trigger pore-collapse. Left: the backbone atoms of cavity-lining residues from two orthogonally opposed Shaker_KD_ subunits (B, D) bound to Cs1 (toxin molecule not shown) are depicted as sticks. Pore ions (tan) and oxygen atoms of cavity water molecules (transparent blue – subunit B, yellow – subunit D) are rendered as spheres. At t~100ns a flip of the carbonyl at Gly444 facilitates the invasion of water molecules from the peripheral pocket in subunit B into the pore, and displacement of the ion bound at S2. This is followed closely by collapse of the pore, and breach of the aromatic cuff barrier in subunit D, enabling flow of water molecules behind the SF into the intracellular vestibule of the pore by the rout depicted in Fig S4F. Right: top view of the pore region demonstrating the asymmetric collapse of the channel pore.

**Figure S5.**
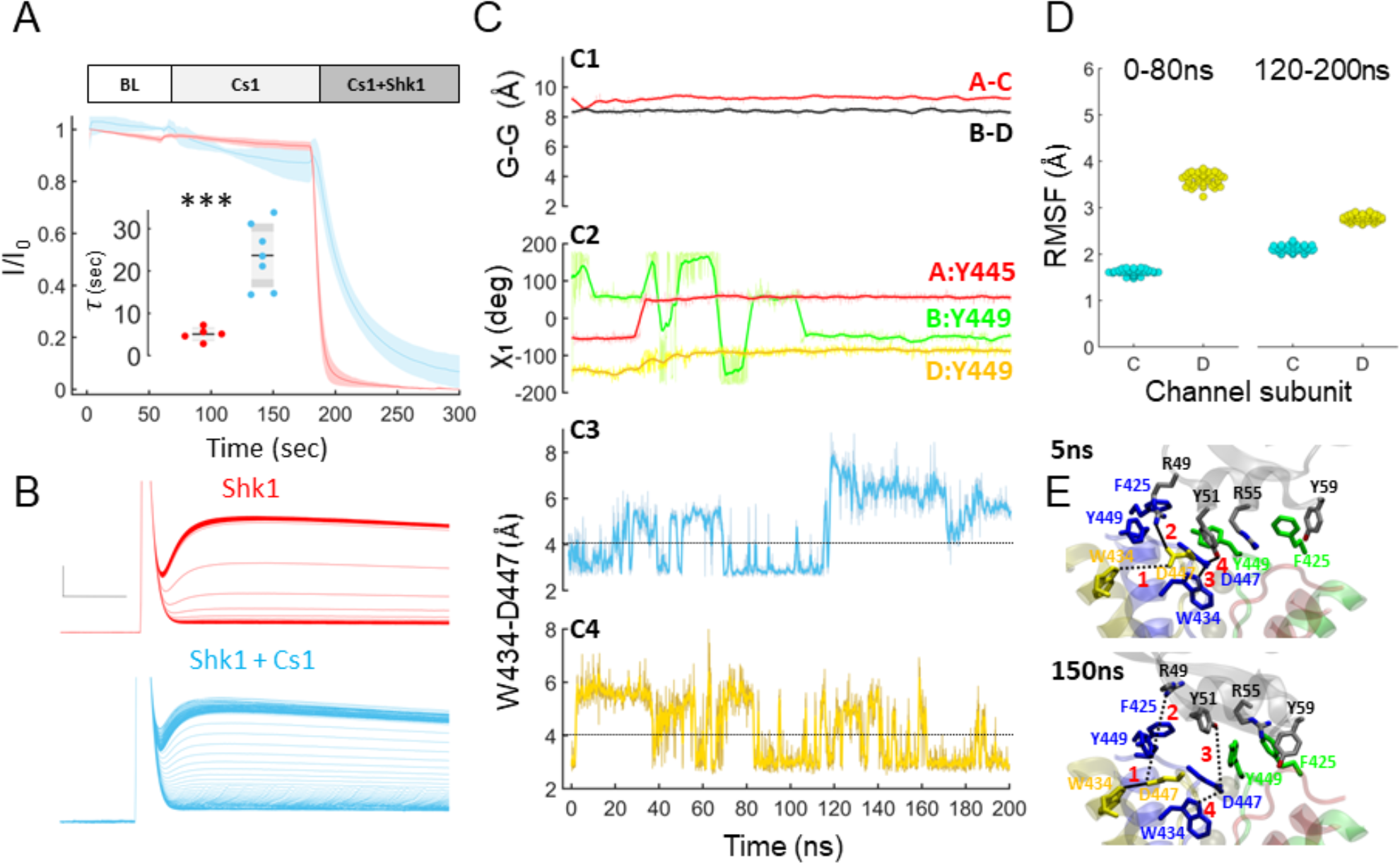
A silent binding mode of Conkunitzin-S1 to a non-inactivating Shaker_KD_ mutant. **A**, Kinetics of Shaker_KD_ block by ShK alone or in the presence of Cs1 at a concentration ratio of 10nM:200nM. See Fig 5A for details. **B**, Raw K^+^ currents recorded at 2s intervals following bath application of 10nM ShK (top) or 10nM ShK followed by 200nM Cs1 (bottom). Scale bars are 10ms/1μA (ShK) and 10ms/0.5μA (ShK^+^Cs1). **C**, Conformational dynamics observed during a 200ns MD production run of the Shaker_KD_ T449Y – Cs1 complex. C1, Cross-subunit distance between the Cα atoms of Gly444 residues from chains A and C (red) or B and D (black) revealing a symmetric pore conformation throughout the trajectory. C2, conformation of the aromatic rings of Y445/Y449 probed by the χ_1_ dihedral angle. B:Y449 (green) is undergoing considerable rearrangements that enable it to engage toxin residues Y51, R55 and Y59 (see movie S5B). A flip of A:Y445 at t~30ns (red) is followed by a rearrangement of D:Y449 (yellow) that stabilizes the aromatic cuff (see movie S5A for details). C3,4 - Dynamics of the DW gate in subunits C (C3) and D (C4). The dashed line marks a 4Å cutoff distance typical of a weak H-bond, applied to score the WD gate as either “open” or “closed”. During the initial 80ns, C:W434-D447 is closed 66.4% of the time, while in chain D is closed only 15% of the time. This pattern is reversed following the conformational rearrangements described above, and at the last 80ns of the run C: W434-D447 is constantly open, while D:W434-D447 is closed 76% of the time. **D**, The difference in thermal motion of water molecules residing at the peripheral pockets of subunits C (cyan) and D (yellow) is diminished following the conformational rearrangement. Each point represents the root mean square fluctuations of pocket waters computed over a 2ns window. The mean RMSF values during the initial 80ns were 1.62±0.05 and 3.61±0.13 for chains C and D, respectively, while during the last 80ns the mean RMSFs were 2.11±0.07 (C) vs. 2.77±0.07 (D). **E**, Shaker_KD_ T449Y-Cs1 complex structure before (5ns, top) and after (150ns, bottom) the structural rearrangement. At t=5ns, the D:W434-D447 bond (1, red) is in an open conformation stabilized via an interaction between D:D447 to Cs1:R49 (2), while the C:W434-D447 bond (3) is in a closed conformation stabilized by a hydrogen bond between C:D447 and Cs1:Y51 (4). At t = 150ns, bonds 2 and 3 are missing, as the sidechains of Cs1:R49 and Cs1:Y51 rearrange to form new cation-*π* and *π*-*π* interactions with the newly introduced Y449 rings (movie S5B).

**Movie S5A.**
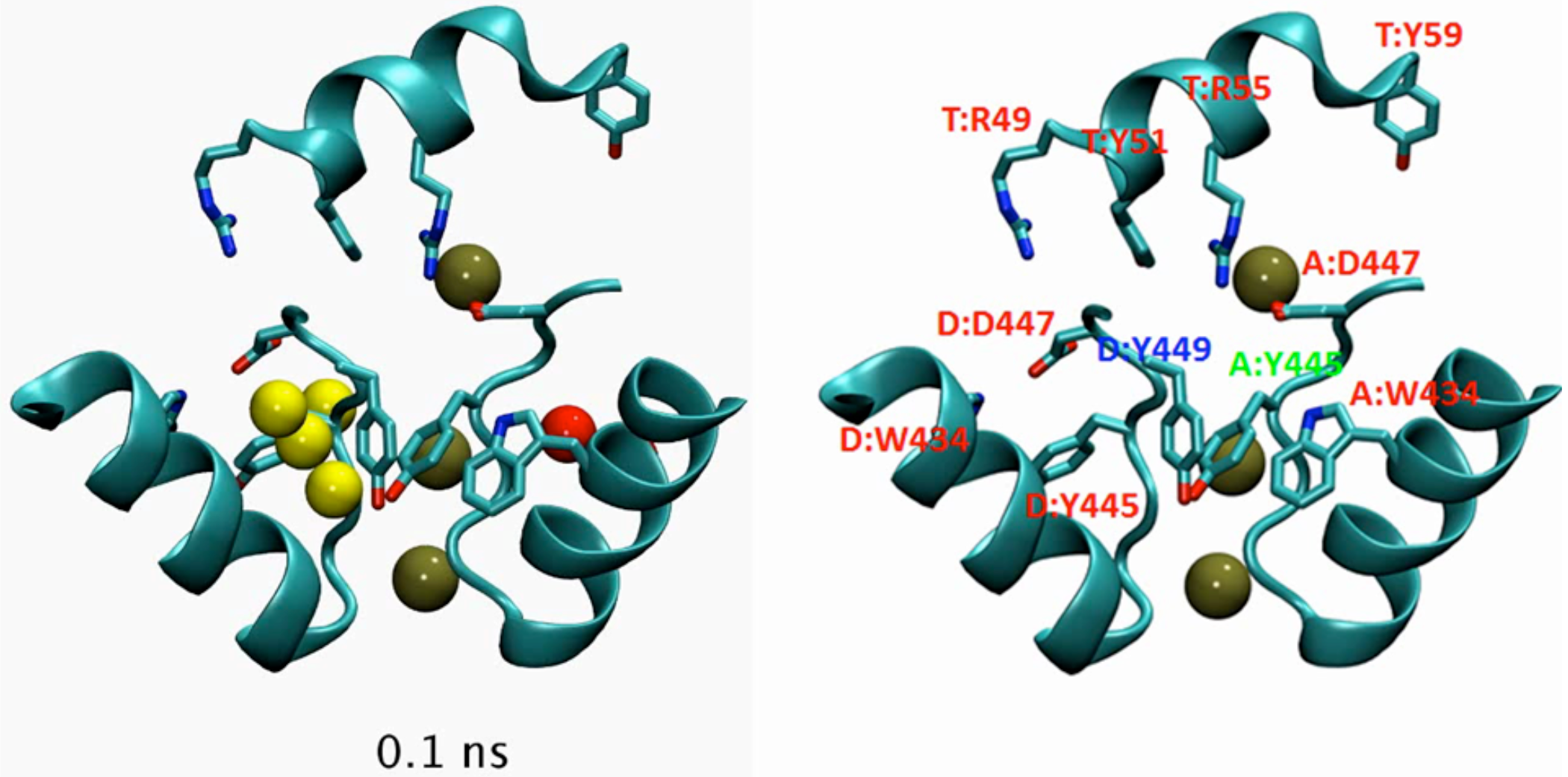
Stabilization of the aromatic cuff barrier by the T449Y substitution. The pore helices of subunits D and A as well as the C-terminal α-helix of Cs1 are shown. The oxygen atoms of water molecules residing at the peripheral cavities D (yellow) and A (red), and K^+^ ions in the pore (tan) are rendered as spheres. Visual legend for the displayed residues is provided at the right. Note the flip of A:Y445 (green legend) at t~30ns, followed by a rearrangement of D:Y449 (blue legend). This rearrangement seem to allow D:Y449 to serve as part of the aromatic cuff-barrier and prevent water movement across the pockets or beneath the barrier as seen in the unmodified channel simulations.

**Movie S5B.**
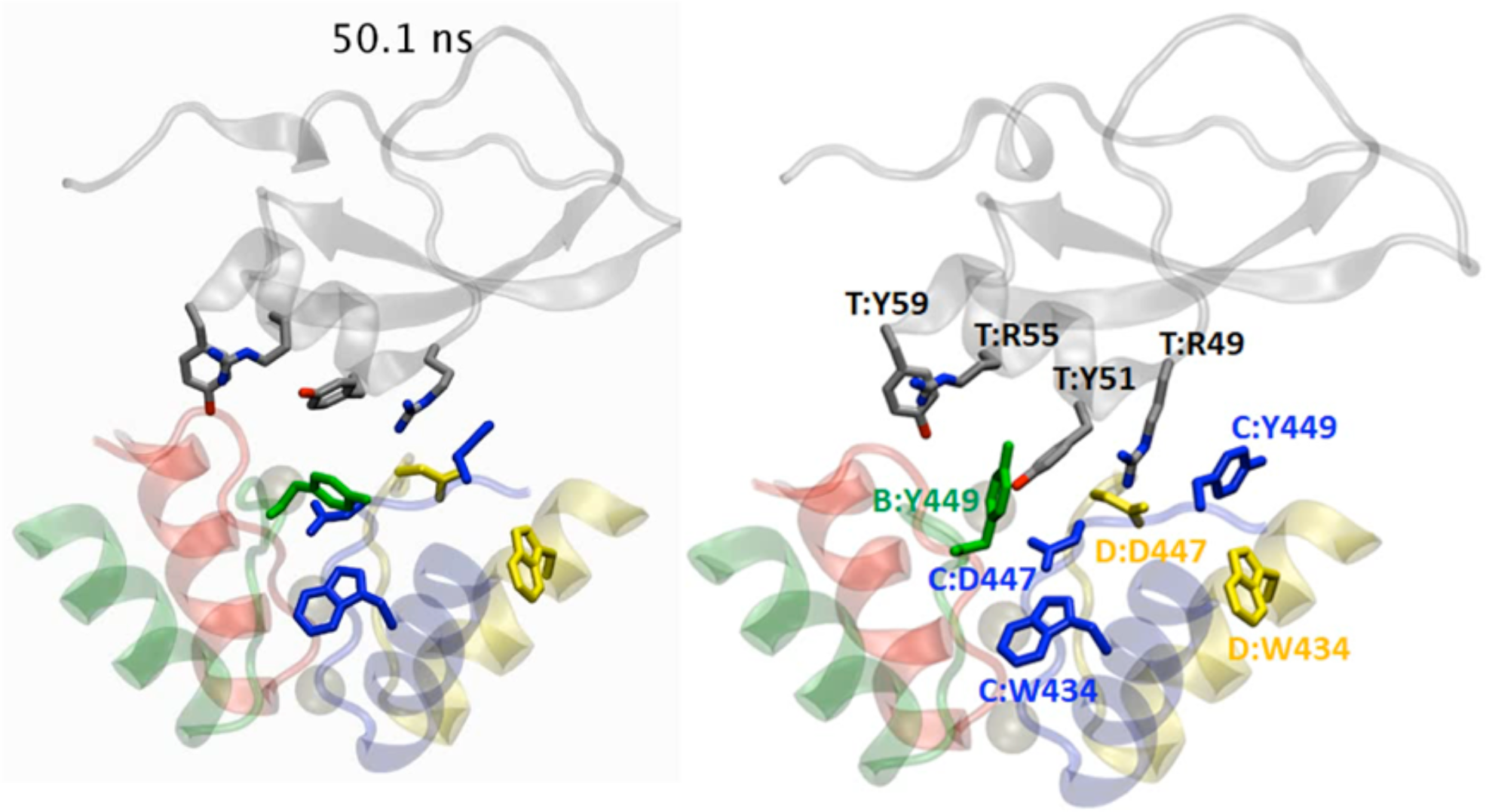
Structural rearrangements within the Shaker_KD_ T449A-Cs1 complex. Channel pore-helices and the toxin are rendered as transparent ribbons, visual legend for the highlighted residues is provided. At t~70ns the sidechain of Cs1:R49 undergoes a rearrangement that abolish its interaction with D:D447. at t~80ns, B:T449 flips into a new conformation that enable it to form novel cation-*π* and *π*-*π* interactions with Y51, R55 and Y59 sidechains of the toxin. In this conformation, Cs1:R55 and Cs1:Y51 are unavailable for interaction with C:D447. The RMSD of the backbone atoms between the first and the last frames displayed is low (2.1Å). Thus the T449Y substitution conserves the overall toxin posture, but abolish the interactions between the toxin to the D447 sidechains by providing an alternative set of interactions. This in turn diminish the toxin effect on the rate of water exchange between the peripheral cavities and the bulk solvent (fig. 5D).

**Figure S6:**
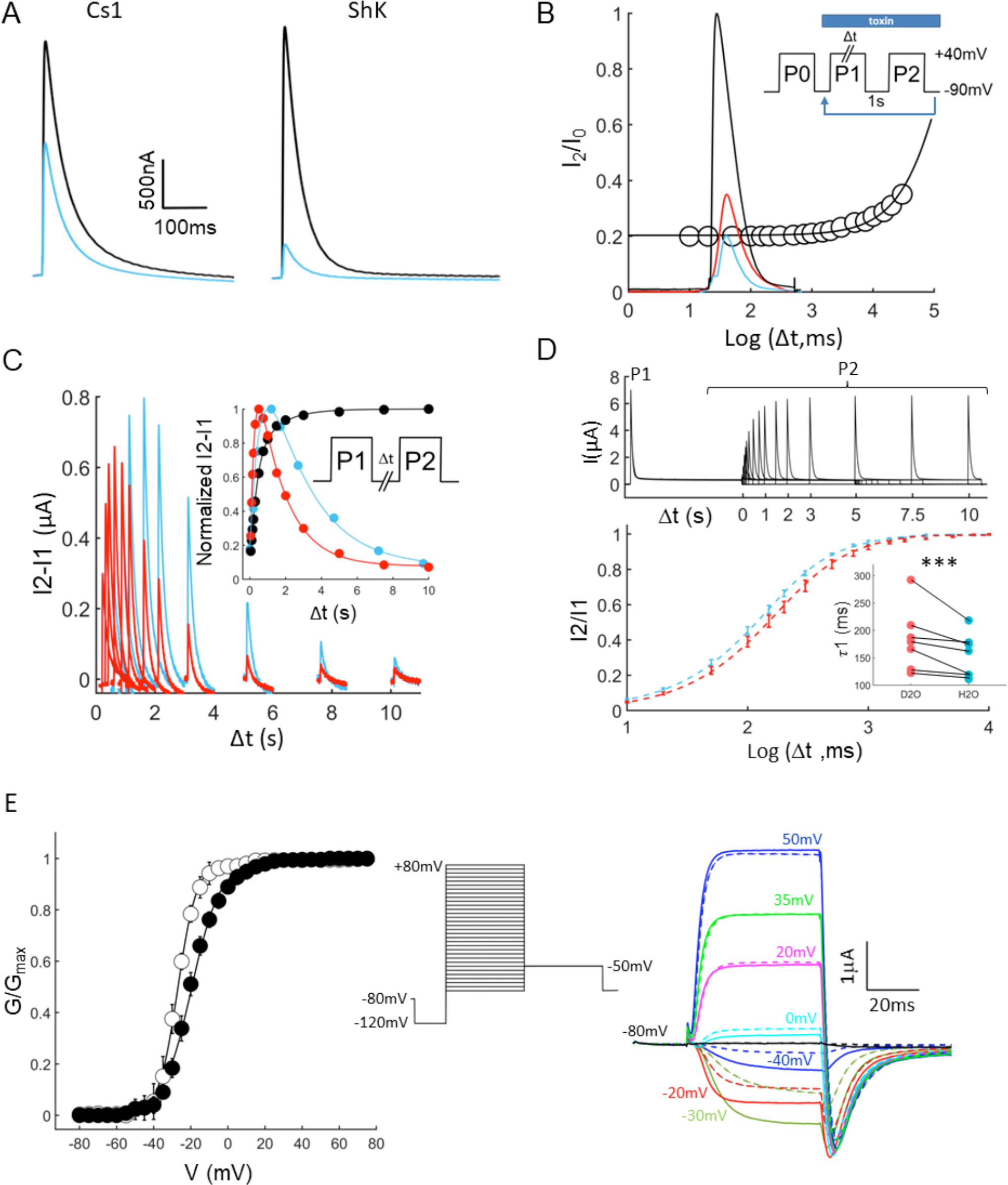
biophysical characterization of the interaction of ShK and Cs1 with Shaker_KD_ M448K. Currents recorded from oocytes expressing Shaker_KD_ M448K during a voltage step to 0mV before (black) and following (blue) application of 2 nM Cs1 (left) or 2nM ShK (right). **B**, Depolarization-induced dissociation of Cs1 from Shaker_KD_ M448K does not require outward K^+^ flux. An oocyte expressing the Channel mutant (black current trace) was exposed to 10nM Cs1, toxin block was allowed to settle for 3 minutes until a steady state amplitude was established (blue trace). A series of depolarizations to +40mV with durations ranging from 0 to 20s was applied, followed by a 1s holding at −90mV to drive recovery from slow inactivation. A second test pulse was then applied and the current amplitude was recorded (open circles). I2 and I0 are the maximal current amplitudes recorded during P2 and P0, respectively. The red trace was recorded at Δt = 20s. Toxin dissociation was best fit using a single exponential with *τ*= 22.5s, while the macroscopic rate of slow inactivation was 35.1 ms, nearly 3 orders of magnitude faster, implying that toxin dissociation takes place mostly in the absence of outward current, setting this phenomenon apart from the trans-enhanced dissociation of classical pore-blockers. **C**, Resettling of toxin block is concentration dependent. An oocyte expressing Shaker_KD_ M448K was exposed to Cs1 at 10 nM (blue) and 50 nM (red). At each toxin concentration, a Two-pulse protocol (fig 6A) was applied to induce reversal and resettling of toxin block. Computed current traces representing the “toxin-released” channel fraction, obtained by subtracting of the traces recorded at P1 from the corresponding traces recorded at P2, are plotted as function of the recovery period. Inset: normalized current amplitude measured during P2 as a function of the recovery period. Black symbols depict channel recovery kinetics in the absence of toxin (*τ*1= 244ms; *τ*2 = 1075ns; A = 0.7), red and blue symbols depict the kinetics in 10nM and 50nM Cs1 respectively, solid lines are best fits to eqn. 4 with *τ*2 = 2866ms (10nM) vs. *τ*2 = 1198 (50nM). **D**, D_2_O decelerates the process of recovery from slow inactivation. A two-pulse protocol (Fig. S6C) was applied to drive channels into slow inactivated state and assay current recovery following subsequent incubations of variable durations (Δt) at holding membrane potential. Top: superposition of currents recorded following recovery periods of 0-10s. Bottom: recovery time course recorded in H_2_O (blue) and D_2_O (red) based solutions. Dashed lines are best fits to a bi-exponential function with *τ*1 = 154±15ms and 183±21ms for H_2_O and D_2_O, respectively, *τ*2 = 828±82, A = 0.8±0.03 (n=7, error bars represent S.E). Inset: *τ*1 values obtained from individual cells in H_2_O (blue) or D_2_O (red), solid lines connect data points derived from the same cell. **E.** Heavy water affect the voltage dependence of Shaker_KD_ activation. Right: current-voltage relationship of the wt channel were acquired in H_2_O (open symbols) or D_2_O (filled symbols) based 90K solutions, and GV relations were extracted from peak tail current amplitudes recorded at a return potential of −50mV. The data were fit with a one component Boltzmann functions with midpoint and slope factor values of V_h_ =-28.3±2.9mV, K_h_= 4.6±0.5mV (H_2_O) and V_h_ =-19.6±1.4mV, K_h_=7.2±1.2 (D_2_O), n = 5. left: superimposed traces recorded from a single oocyte in either H_2_O (solid lines) or D_2_O (dashed lines) based solutions at the indicated test potentials.

**Table S6:**
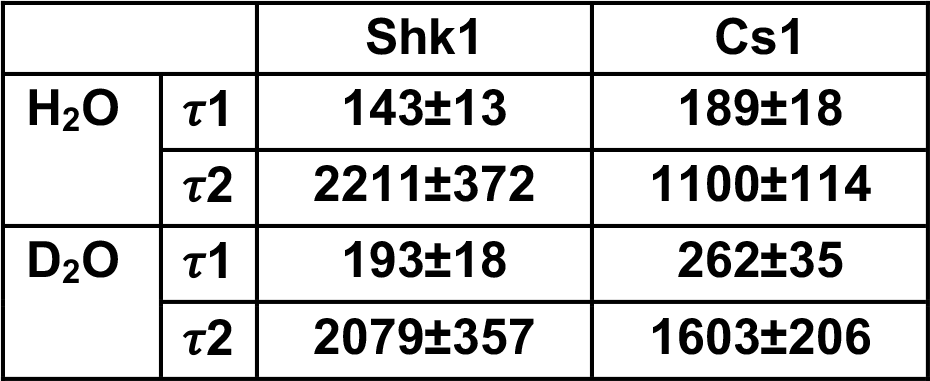
time constants obtained by fitting the toxin re-settling curves presented in Fig 6 B, C using equation 4.

